# A characterization of mitotic and centrosomal defects in a continuum model of Breast Cancer

**DOI:** 10.1101/2024.09.20.614099

**Authors:** Alexsandro dos Santos, Caroline Diorio, Francine Durocher, Sabine Elowe

**Affiliations:** Département de Médecine Moléculaire, Faculté de Médecine, Université Laval, Québec, QC G1V 0A6, Canada; Centre de Recherche sur le Cancer, CHU de Québec-Université Laval, Québec City, QC G1V 4G2, Canada; PROTEO-regroupement québécois de recherche sur la fonction, l’ingénierie et les applications des protéines, Québec, QC G1V 0A6, Canada; Département de médecine sociale et préventive, Faculté de médecine, Université Laval, Québec, Canada; Département de Pédiatrie, Faculté de Médecine, Université Laval et le Centre de recherche sur le cancer de l’Université Laval, Québec, QC G1R 2J6, Canada

## Abstract

Errors in mitosis can contribute to aneuploidy and CIN and play a pivotal role in cancer. So the identification of altered mitotic regulators can contribute to the understanding of the development and progression of breast cancer. In the present study we used an *in vitro* model of disease progression (the MCF10A series of BC continuum) and analyzed the errors of chromosome segregation that occur during the progression of the disease. Our findings indicated that the MCF10A series exhibited several abnormalities in chromosome segregation and its frequency increased with the disease progression. These errors included anaphase lagging chromosomes, micronuclei, nuclear buds, nucleoplasmic bridges, errors of chromosome alignment, and centrosome loss/amplification. Moreover, the presence of centrosome amplification disrupted the proper orientation of the mitotic spindle, resulting in the generation asymmetrical cell lines and aneuploidy in the MCF10A series. Hyper stable kinetochore-microtubule (kt-MT) attachment was also found in premalignant, preinvasive, and invasive cell lines, which can also explain the presence of errors of chromosome alignment. The human transcriptome array also determined possible negative regulators of ciliogenesis that can explain the mechanism of chromosome missegregation that lead to CIN found in the MCF10A series. Collectively, these findings highlight the importance of mitotic defects in the progression of breast cancer.

## Introduction

Aneuploidy was first observed in tumor cells over 100 years ago by the German zoologist Theodor Boveri while studying sea urchin embryos undergoing abnormal mitotic divisions^1^. This feature is a state in which the number of chromosomes in a cell or organism deviates from multiples of the haploid number of chromosomes^2,3^. It may occur by chromosome gains and losses due to chromosome segregation errors, a so called “whole chromosomal” aneuploidy, as well as due to rearrangements of chromosomal parts, often accompanied by their deletion and amplification, that is referred to as a “structural” or “segmental” aneuploidy. Frequently, a combination of both structural and numerical chromosomal changes can be found in cancer cells (composite aneuploidy)^4^.

Aneuploidy is often associated with a more complex phenotype named chromosomal instability (CIN)^5–7^. It is important to note that both terms (aneuploidy and CIN) are not synonymous: aneuploidy describes the state of having an abnormal chromosome number (the “state” of the karyotype), and the second one refers to an elevated rate of chromosome gain or loss (the “rate” of karyotypic change) ^1,8,9^. Aneuploidy therefore may be a possible ‘gateway’ to increasingly elevated CIN. Similarly, CIN may lead to aneuploidy, as the effects of CIN will invariably lead to structural and/or numerical aneuploidy^9^.

CIN is a hallmark of human cancers and there is a growing evidence that it is correlated to tumor development^1,10,11^ and poor prognosis in patients^12–15^. Furthermore, CIN can also drive tumor heterogeneity, where aneuploidy can have a causal role^16^, and contributes to clonal evolution, where increased chromosome mis-segregation events lead to the generation of cells with proliferative advantages, and drug resistance^17–19^.

Breast cancer (BC) is a highly complex and heterogeneous disease with some cases being associated with slow growth and excellent prognosis whilst other tumors exhibit a highly aggressive clinical course^20^. This disease ranks as the fifth cause of death from cancer overall, is the most frequent cause of cancer death in women in less developed regions (14.3% of total), and the second cause of cancer death in more developed countries (15.4%) after lung cancer^21^. Recent GLOBOCAN (Global Cancer Statistics) data produced by the IARC (International Agency for Research on Cancer) from 185 countries estimated that 2.26 million women were diagnosed with BC, and 684,000 died from this disease worldwide in 2020^22^.

There are several models that explain the BC progression. The linear model of disease progression states that BC progresses stepwise through different stages, which it initiates as the premalignant stage of atypical ductal hyperplasia (ADH), progresses into the preinvasive stage of ductal carcinoma *in situ* (DCIS), and culminates in the potentially lethal stage of invasive ductal carcinoma (IDC)^23^. In this model, ADH and DCIS are nonobligate precursors of IDC. The nonlinear (or branched) model states that DCIS is an obligatory progenitor of IDC, yet different grades of DCIS progress to corresponding grades of IDC^24^. Finally, the ‘parallel’ model of progression of DCIS and IDC hypothesizes that DCIS and IDC diverge from a common progenitor cell and progress independently through different grades in parallel^25^. To study breast tumor progression, a powerful cell model (the MCF10A series) was developed. Briefly, the basis of the model is a human cell line MCF10A (Figure 1A). This cell line is an immortal human breast epithelial cell line originating from subcutaneous mastectomy of a breast with benign fibrocystic disease from a 36-year-old woman^26^. Transfection of MCF10A cells with the mutated human T-24 HRAS gene resulted in cells with a transformed phenotype (MCF10AneoT) which have lost anchorage dependence *in vitro*^26^. Then MCF10AneoT was injected into immunodeficient nude mice to develop MCF10AT1^27^. After two successive trocar passages initiated from a xenograft lesion formed by MCF10AT1, MCF10DCIS.com was obtained^28^. Next, after multiple passages of MCF10AT1 into nude mice, MCF10ACA1a was generated^29^. MCF10A (normal breast), MCF10AT1 (premalignant cell line), MCF10DCIS.com (preinvasive cell line), and MCF10CA1a (invasive cell line) recapitulate successive steps in BC progression like normal breast, ADH, DCIS and IDC, respectively. The continuum represents an isogenic model of disease progression and has been widely used to characterize the genetic drivers of breast malignancy^30,31^.

**Figure 1.**
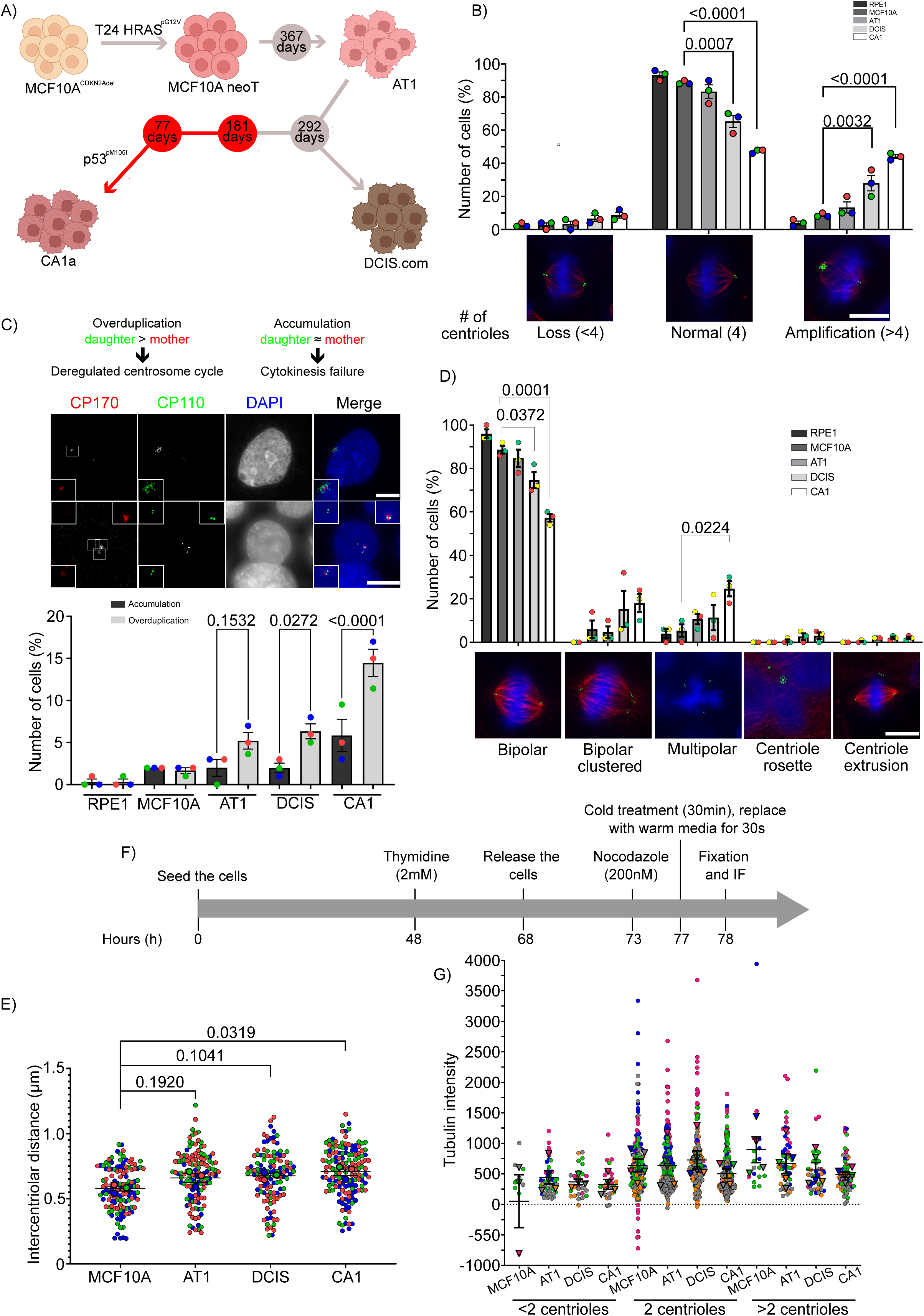
MCF10A series presented centriole loss and amplification and these characteristics increased with disease progression. A) Diagrammatic representation of the generation of the MCF10A series. Numbers inside the circles correspond to the number of days for which cell line was grown in vivo before reinplantation. B) Graph showing the percentage of cells presenting centrosome loss (<4 centrioles), normal centrosomes (4 centrioles) and centrosome amplification (>4 centrioles) (n=150 cells). Representative images of cells showing centrosome loss (image on the left), normal centrosomes (middle) and centrosome amplification (image on the right). Cells were stained with anti-α-tubulin (red), anti-CP110 (green), and DAPI (blue). C) Mechanism of centrosome amplification in MCF10 series. Cells were stained with anti-CP110 (green), anti-CEP170 (red), and DAPI (blue) to distinguish between centriole accumulation or overamplification. The graph shows the percentage of cells with the indicated phenotypes (overduplication or accumulation). (n=300 cells). D) Spindle organization in MCF10A series. Graph showing the different mechanisms present in MCF10A series to cope with centriole amplification (bipolar spindle, bipolar clustered, multipolar, centriole rosette, and centriole extrusion). Representative images of cells presenting the phenotypes described. Cells were stained with anti-α-tubulin (red), anti-CP110 (green), and DAPI (blue). (n=150 cells). E) Intercentriolar distance (μm) in MCF10A series. F-G) Microtubule nucleation assay. F) Timeline of the microtubule regrowth assay. G) Quantification of MT regrowth in MCF10A series. <2: cells with less than 2 centrioles per foci; 2: cells with 2 centrioles per foci; >2: cells with more than 2 centrioles per foci. All statistical tests of comparative data were performed using one-way ANOVA for differential comparison between more than two groups. Data are shown as mean ± SEM. P < 0.05 were considered statistically significant. n = 3 biological replicates (except microtubule nucleation assay with n = 5 biological replicates). Scale bar: 10µm

Several studies have shown that genes implicated in aneuploidy arising from chromosome missegregation errors and CIN are significant predictors of clinical outcome in several BC datasets^32–35^. For example, it is estimated that most breast tumors (about 75%) present some degree of CIN^12^. The mechanisms that lead to CIN are poorly understood but mostly reflect dysfunctional chromosome segregation during mitosis^1,2,36^. These mechanisms reduce mitotic^37^ fidelity and include defects in chromosome cohesion, the spindle assembly checkpoint (SAC), centrosome copy number, kinetochore– microtubule attachment dynamics, and cell-cycle regulation^2^. Additionally, our group identified a specific gene signature that could represent potential biomarkers for each stage of BC progression (ADH, DCIS and IDC) and which could be used by the clinicians in early diagnosis and treatment intervention^38^.

Interestingly, our group showed that progression to IDC was associated with significant changes in the expression of mitotically relevant genes. Recently, using a list of mitotic genes generated through curation of mitotically relevant GO (gene ontology) annotation terms, we queried a patient-derived BC continuum gene expression dataset and identified cytoskeleton-associated protein 2 (CKAP2) as an important mitotic regulator in invasive BC^39^. Thus, we hypothesize that mitotic defects that contribute to CIN play a critical role in both the development and progression of BC. In this study, using an *in vitro* model of disease progression (the MCF10A series of BC continuum), we analyzed and described the missegregation defects that occur during the progression of the disease and lead to CIN. Our results showed that the MCF10A series presented an array of chromosome segregation defects, including lagging chromosomes, micronuclei, nuclear buds, nucleoplasmic bridges, errors of chromosome alignment, centrosome loss/amplification, which increased with the disease progression. Additionally, the time spent in mitosis increased with the disease progression due to the increasing errors of chromosome alignment. Furthermore, centrosome amplification disrupted mitotic spindle orientation, which can lead to the formation of asymmetrical cell lines and aneuploidy in the MCF10A series. Premalignant, preinvasive, and invasive cell lines presented hyper stable kinetochore-microtubule (kt-MT) attachment, which can also explain the presence of errors of chromosome alignment in those cell lines. Moreover, there is a loss of primary cilia according to BC progression. We also identify possible negative regulators of ciliogenesis that can explain the mechanism of chromosome missegregation that lead to CIN found in the MCF10A series. Taken together, these results show the importance of mitotic defects in the progression of BC.

## Material and methods

### Cell lines

MCF10A and MCF10AT1 (hereafter AT1) cells were cultured in Dulbecco’s Modified Eagle Medium (DMEM) F-12 media (Wisent Inc, Québec, CA) supplemented with 5% horse serum (HS) (Sigma Aldrich, Oakville, CA), 10 mM HEPES (4-(2-hydroxyethyl)-1-piperazineethanesulfonic acid) (Fisher BioReagents, CA), 20 ng/ml epidermal growth factor (EGF) (GIBCO, Carlsbad, CA), 0.01 mg/ml insulin (Wisent Inc, Québec, CA) and 500 ng/ml hydrocortisone (Sigma Aldrich, Oakville, CA). MCF10DCIS.com (hereafter DCIS) and MCF10CA1a (hereafter CA1) were cultured in DMEM F-12 supplemented with 5% HS and 10mM HEPES.

hTERT-RPE1 (hereafter RPE1) is a chromosomally stable diploid retinal pigment epithelium cell line immortalized by constitutive expression of human telomerase^40^. RPE1 was cultured in Dulbecco’s Modified Eagle Medium (DMEM) F-12 media (Wisent Inc, Québec, CA), supplemented with 10% fetal calf serum (Sigma Aldrich, Oakville, CA).

All media were supplemented with 100 U/mL penicillin and 100 μg/mL streptomycin (HyClone™ Penicillin Streptomycin 100X Solution, Thermo Fisher Scientific).

### Immunostaining

Cells grown on polylysine-coated coverslips (500 µg/mL poly-L-lysine), and after fixation with specific methods, were blocked in 3% BSA (bovine serum albumin) in 1X PBS (phosphate-buffered saline) for 30 min. Cells were then incubated with primary antibody for 1 hour at room temperature and washed three times with 1X PBS solution. Secondary antibody incubation was performed for 1 hour at room temperature before final washing and mounting on microscopy slides.

### Manual scoring of micronuclei (MNi), nuclear buds (NB) and nucleoplasmic bridges (NPB)

Asynchronous DAPI (4′,6-diamidino-2-phenylindole)-stained MCF10A series were analyzed. Images were manually scored for micronuclei (MNi), nuclear buds (NBs) and nucleoplasmic bridges (NPBs). The criteria used to identify MNi, NBs and NPBs was according to Fenech^41^. Briefly, micronuclei are morphologically identical to but smaller than nuclei (their diameter should not exceed 1/3 that of the primary nucleus); they are not linked or connected to the main nuclei; they may touch but not overlap the main nuclei; they have the DAPI staining intensity as the main nuclei but occasionally staining may be more intense. NPB is a continuous DNA-containing structure linking the nuclei in a binucleated cell; its width may vary considerably but usually does not exceed 1/4th of the diameter of the nuclei within the cell; it should also have the same DAPI staining characteristics as the main nuclei. NPBs are similar to MNi (the exception is that they are connected with the nucleus via a bridge that can be slightly narrower than the diameter of the bud or by a much thinner bridge depending on the stage of the extrusion process); usually they have the same DAPI staining intensity as MNi.

### Lagging chromosomes

In this experiment, 300,000-375,000 cells were seeded on poly-L-lysine-coated coverslips (500 µg/mL poly-L-lysine). After 48h, cells were incubated with RO-3306 (4 µM) for 16hours. Then RO-3306 was washed out and cells were released for 1.5 hour, in order to have cells arrested in anaphase. After that, cells were fixed with PTEMF buffer (20 mM PIPES pH 6.9, 0.2% Triton X-100, 10 mM EGTA, 1 mM MgCl2, and 4% formaldehyde) and then immunostained. Primary antibodies used were CREST anti-centromere serum (1/1000 dilution, HCT-0100, human monoclonal, Immunovision) and anti-α-tubulin (1/1000 dilution, DM1A, sc-32293, mouse monoclonal, Santa Cruz). Hoechst 33342 (Thermo Scientific) was used at 1 mg/ml.

Lagging chromosomes in anaphase were scored manually and defined as any DAPI-positive material between the two chromosome masses (with or without centromere marker), but distinguishably separated from them (excluding NPBs)^42^.

### Microtubule regrowth

Microtubule regrowth assay was performed as described previously^43^ with some modifications. Briefly, 300,000-375,000 cells were seeded on poly-L-lysine-coated coverslips. One day after, cells were arrested in S phase by the addition of 2 mM thymidine for 20 hours. After that, thymidine was washed out and cells were released for 5 hours. Cells were subsequently incubated with 200 nM nocodazole for 5 hours. Afterwards, cells were washed 3 times with cold media and placed on ice bath for 30 minutes. Regrowth of microtubule was induced after replacing the cold media with prewarmed media at 37 °C. After 30 seconds, media was removed and cells were fixed with PTEMF and then immunostained (see the immunostaining section for details). Primary antibodies used were anti-centrin-2 (1/1000 dilution, clone 20H5, mouse monoclonal, Millipore) and anti-α-tubulin (1/1000 dilution, DM1A, sc-32293, mouse monoclonal, Santa Cruz). Hoechst 33342 (Thermo Scientific) was used at 1 µg/ml.

The quantification of microtubule regrowth was done as previously published^44,45^. Briefly, a 5-μm circle was drawn in order to position the centrioles in the center, then the channel was switched to the one showing the microtubule staining and the intensity of the selected region was measured. The intensity of a microtubule-free background region of the same area was subtracted from the sample data to obtain the background-corrected intensity of the microtubules.

### Centriole counts

Cells arrested in metaphase (as described in cold stable assay) were used for this analysis. After that, cells were fixed with cold methanol for 10 minutes at −20°C and then immunostained. In order to count the number of centrioles with no bias^46^, two antibodies were used: anti-centrin-2 (1/1000 dilution, clone 20H5, mouse monoclonal, Millipore) and anti-CP110 (1/500 dilution, 12780-1-AP, rabbit polyclonal, Proteintech). Hoechst 33342 (Thermo Scientific) was also used at 1 µg/ml. Both centrin-2 and CP110 localize to the distal tip of centrioles and typically appear as four foci in mitotic cells^47^. Centrioles were counted manually and only centrioles labeled with both markers were considered. Cells were scored as centriole loss (< 4 centrioles), normal (4 centrioles) and centriole amplification (> 4 centrioles).

### Cold stability assay

Here, 300,000-375,000 cells were seeded on poly-L-lysine-coated coverslips (500 µg/mL poly-L-lysine). After 48 hours, cells were incubated with the CDK1 inhibitor RO-3306 (4 µM) for 16 hours. Cells were then washed and released for 1 hour into media containing MG-132 (carbobenzoxy-l-leucyl-l-leucyl-l-leucinal) (10 µM), resulting in a metaphase-arrested population. After that, cells were incubated for 20 and 30 minutes on ice, then fixed for 10 minutes at room temperature with PTEMF. Finally, fixed cells were immunostained as described. Primary antibodies used for this experiment included anti-α-tubulin (1/1000 dilution, DM1A, sc-32293, mouse monoclonal, Santa Cruz) and anti-CP110 (1/500 dilution, 12780-1-AP, rabbit polyclonal, Proteintech). Hoechst 33342 (Thermo Scientific) was used at 1 µg/ml.

### Primary cilia

For the induction of primary cilia, serum starvation was used^48^. Briefly, cells were seeded on coverslips and cultured in growth media until confluency. After that, old media was changed for a free-serum media. Cells were fixed at indicated times (5, 7, and 10 days after serum starvation) in 3.7% paraformaldehyde (PFA) for 20 minutes at room temperature (RT) followed by permeabilization with 0.3% Triton X-100 in PBS for 10 minutes at RT, and then immunostained as described. Primary antibodies used included anti-acetylated α-tubulin (1/1000 dilution, clone 6-11B-1, mouse monoclonal, Sigma) and anti-CP110 (1/500 dilution, 12780-1-AP, rabbit polyclonal, Proteintech). Hoechst 33342 (Thermo Scientific) was used at 1 µg/ml.

### Mitotic spindle morphology analyses

Here, 300,000-375,000 cells were seeded on poly-L-lysine-coated coverslips (500 µg/mL poly-L-lysine). After 48 hours, cells were incubated with the CDK1 inhibitor RO-3306 (4 µM) for 16 hours. Cells were then washed and released for 1 hour into media containing MG-132 (10 uM). After that, cells were fixed with methanol and immunostained using anti-α-tubulin (1/1000 dilution, DM1A, sc-32293, mouse monoclonal, Santa Cruz) and anti-CP110 (1/500 dilution, 12780-1-AP, rabbit polyclonal, Proteintech) as primary antibodies to stain mitotic spindle and centrioles, respectively. Moreover, Hoechst 33342 (Thermo Scientific) was used at 1 µg/ml.

Measurements were assessed using ImageJ (Fiji package). Spindle length (pole-to-pole distance) was defined as the pole-to-pole distance between two centrioles foci. Half-spindle length was defined as the distance from the center of the centrioles foci to the center of the chromatin at metaphase^49^. Half-spindle ratio was defined as the ratio of the two half-spindle lengths^50^. Chromosome congression index was defined as the ratio between width and length of metaphase plate (Width/Length) stained by DAPI^51,52^. An ImageJ macro was used to calculate, by inverse trigonometric function, the mitotic spindle angle^53^. These measurements derive from the two spindle pole coordinates. Negative angles or angles above 90° were transformed to the first quadrant. Angular histograms were plotted using a custom R script kindly provided by Dr. Elena Scarpa^54^. Intercentriolar distance, which corresponds to the distance between two centrioles in a pair, was measured from the center of one centriole to the center of its respective centriole pair.

Centriole markers (centrin-2 or CP110) were used to classify mitotic spindle morphology as bipolar, bipolar clustered and multipolar^55^. Bipolar spindle was defined as a spindle that presented two opposite poles with two centrioles per pole. Bipolar clustered corresponds to a spindle with two opposite poles and, at least, 3 centrioles clustered in one pole. Spindles with three or more poles were considered as multipolar. Centrioles can also assemble in a way that resembles a ‘rosette’ arrangement – in this arrangement, a mother centriole is surrounded by numerous daughter centrioles^56^. Centrioles can also be eliminated/degraded by a mechanism called centriole extrusion where they are degraded/extruded from the cells^57^.

### Time-lapse observation of live cells and confocal microscopy

Time-lapse observations and confocal microscopy were performed using an Olympus IX80 inverted confocal microscope equipped with a WaveFX-Borealin-SC Yokagawa spinning disc (Quorum Technologies) and an Orca Flash4.0 camera (Hamamatsu).

For confocal microscopy, images shown represent Z-projection of 20 independent acquisitions, with a distance between planes of 0.2 μm.

For live cell imaging, cells were seeded in an 8-well chamber slides (Thermo Scientific Nunc Lab-Tek) at a density of 2 x 10^4^ cells per well. One day after, cells were incubated with 2 mM thymidine for 24 hours, then released for 2 hours. Two hours before imaging, cells were incubated with 0.25 µM siR-Hoechst (SiR-DNA; Spirochrome). Time-lapse imaging was acquired at 20X objective every 3 minutes at 37°C, 5% CO_2_ for 12 hours.

Images from time lapse experiments and confocal microscopy shown in the same figure have been identically scaled. Imagine processing was performed using the plugin QuickFigures from ImageJ^58^.

### Proliferation assays

MCF10A series and hTERT RPE-1 were seeded in duplicates onto 35 mm plates (25,000 cells per plate). After attachment (usually 2 days), cells were treated with the PLK1 inhibitor centrinone (200 nM) to deplete centrosomes or DMSO as control. On days 1, 2, 3, 4, 5, and 7 of centrinone treatment, cells were rinsed with cold PBS and trypsinized. Once trypsin was neutralized, cells were gently mixed, fixed in 1 ml final volume of 3.7% formaldehyde, and counted on the hemocytometer. Media containing fresh centrinone replaced old media every three days.

### Microarray dataset and identification of differentially expressed Genes

In this study, gene expression data was obtained from our previous human transcriptome array (HTA) performed in MCF10A cell line subtypes (MCF10A, AT1, DCIS, and CA1)^38^. Differentially expressed genes (DEGs) with a fold change (FC) ≥ |2.0| and a P value cutoff of < 0.05 were considered as statistically significant.

### Gene function and pathway enrichment analysis by Metascape

Gene function and enrichment analyses for DEGs were performed using Metascape (http://metascape.org/, accessed on 15 September 2022)^59^. Pathway analysis was performed using Reactome gene sets, canonical pathways, BioCarta gene sets, Gene Ontology (GO) biological processes, Hallmark gene sets, and Kyoto Encyclopedia of Genes and Genomes (KEGG); functional analysis was performed using GO molecular functions; and structural complex analysis was conducted using GO cellular components, KEGG structural complex, and CORUM (comprehensive resource of mammalian) protein complex. Terms with a P value < 0.05, a minimum count of 3, and an enrichment factor of > 1.5 were collected and grouped into clusters based on their membership similarities. The most significant term within a cluster was selected as the one representing the cluster.

### Integration of protein–protein interaction (PPI) network and module analysis

Protein-protein interaction (PPI) networks of intersection DEGs were extracted from Metascape and visualized in Cytoscape (version 3.9.1)^60^. The network, which was created taking into consideration only physical interactions in STRING (physical score > 0.132) and BioGrid, contains the subset of proteins that form physical interactions with at least one other member in the gene list. The Molecular Complex Detection (MCODE) was applied to identify densely connected network components and most significant module in PPI network.

### Statistical Analysis

Statistical analysis was performed with GraphPad PRISM software version 9.5.0 (San Diego, CA, USA). All assays were performed in triplicates (exception growth curve done in duplicates) and repeated at least three times. Statistical analysis for the comparisons of means culture were done using one-way or two-way ANOVA. Angular histograms were compared using Kolmogorov-Smirnov test. P values smaller than 0.05 are considered significant.

## Results

### MCF10A series presented centriole loss and amplification and these characteristics increased with disease progression

Centrosome amplification (CA) is a common feature in cancer^46,61,62^ and correlates with more aggressive clinical phenotype and worse patient outcomes^61,63,64^. To determine the frequency of CA in BC progression, we analyzed centrosomes by immunofluorescence (IF) staining using two centriole markers (centrin-2 and CP110). The use of both markers facilitated counting in a non-biased manner the number of centrioles, and only centrioles doubly labelled were counted ^46^. There was a significant decrease in the frequency of cells presenting 4 centrioles as the disease progresses (MCF10A, AT1, DCIS, and CA1 with a mean of 88.67, 83.33, 65.33, and 47.33%, respectively) (Fig 1B). Moreover, there was a significant increase of centrosome amplification according to disease progression (MCF10A, AT1, DCIS, and CA1 presented mean of 8.67, 13.33, 28.00, 44.00%, respectively). In both cells with normal centriole number and CA, there was no difference in mean percentage of MCF10A cells when compared to the RPE1. Interestingly, there was also centriole loss (cells with < 4 centrioles) in the series. The frequency was low, although there were no differences in the mean percentage among steps of BC progression (the mean percentage found in MCF10A, AT1, DCIS, and CA1 cells was 2.67, 3.33, 6.67, and 8.70%, respectively). These results showed that CA amplification is a common feature in BC, it is present before the development of aggressive tumors, and correlates with the progression of the disease, in agreement with other BC studies^61,62,65–67^.

CA can be a result of two major mechanisms: (1) centriole overduplication and (2) centriole accumulation^68–71^. Centriole overduplication leads to the excess of daughter centrioles surrounding one or two maternal centrioles. Additionally, centrosome accumulation is characterized by the presence of multiple sets of normally replicated centrosomes, each set composed of a mother-daughter couple. The difference between these two mechanisms is that the overduplication is caused by deregulation of centrosome duplication while accumulation results from events that are not related to the centrosome duplication process, such as cytokinesis failure or mitotic slippage^72^. Hence, we decided to evaluate the contribution of the two main mechanisms of CA in the series (Figure 1C). To distinguish between these two mechanisms, double staining IF using centrin-2 and CEP170 was performed^73,74^. Centrin-2 is present in both mother and daughter centrioles^75^ and CEP170 associates exclusively with subdistal appendages of mature mother centrioles ^74^. When centrin-2 co-stains with CEP170, that centriole is considered mother centriole. Otherwise, a centriole stained only with centrin-2 is considered a daughter. Therefore, centriole overduplication is the predominant mechanism leading to CA when the number of daughter centrioles is greater than the number of mother centrioles. Nevertheless, centriole accumulation is the predominant mechanism when the number of mother and daughter centrioles is balanced. The graph in Figure 1C shows the quantification of overduplication and accumulation in the series. There was no difference between centriole overduplication and accumulation found in MCF10A cells (mean percentage of overduplication and accumulation was 1.67 and 2.00%, respectively). However, for AT1, DCIS, and CA1 cells, most of the centriole amplification originated from centriole overduplication. For AT1 cells, the mean percentage of overduplication and accumulation was 5.21 and 2.00%, respectively (p = 0.1532). For DCIS cells, the mean percentage for overduplication and accumulation was 6.33 and 1.98% (p = 0.0272), respectively. For CA1 cells, the mean percentage for overduplication and accumulation was 14.47 and 5.84 (p < 0.0001), respectively. These results show that centriole overduplication is the predominant mechanism driving the centriole amplification in AT1, DCIS and CA1 cells.

Centrosome amplification (generated either by centriole overduplication or cytokinesis failure) has been proposed to generate multipolar spindle which leads to asymmetric chromosome segregation, which will result in increased aneuploidy and CIN^66,76,77^. However, if the spindle multipolarity is not corrected it will invariably lead to cell death^78^. In order to avoid these mitotic pitfalls, cells display a number of coping mechanisms (including centriole clustering, centriole rosette and centriole extrusion)^56,79^. We decided to quantify which coping mechanisms are present in order to avoid cell death (Figure 1D). As depicted in the graph, there is a decrease in the frequency of bipolar spindle formation with disease progression and a corresponding increase in the percentage of cells with multipolar spindles and bipolar spindles with clustered centrosomes (although there was no significant difference). Moreover, centriole extrusion and centriole rosettes were also found, however, in a very low frequency (less than 5% for all steps of the BC progression). These results show that BC cells use a wide range of mechanisms to cope with centriole amplification to avoid multipolar spindle.

In normal cells, the centriole is duplicated only once per cell cycle during S phase. The newly formed centrioles remain engaged with their mother centrioles until the end of mitosis, when they disengage. This step licenses a new round of centriole duplication in the next cell cycle^80^. If disengagement, defined as a loss of orthogonal orientation between centrioles, occurs prematurely it can also lead to the formation of multipolar spindles independent of centrosome amplification^81^. The distance between two centrioles in a centrosome is a readout for the centriole disengagement, and distances greater than 0.65 μm indicate that centrioles are either disoriented or moving apart (disengaging)^82^. As depicted in the graph (Figure 1E), when we evaluated the intercentriolar distance, we found no significant difference in the average distance between centrioles in AT1 (0.70 µM ± 0.01, p = 0.0192) and DCIS cells (0.67 µM ± 0.02, p = 0.1014) when compared to MCF10A cells (0.56 µM ± 0.01). However, there was a significative difference in the intercentriolar distance mean in CA1 cells (0.74 µM ± 0.08, p = 0.0319) when compared to MCF10A cells. These results show that centrioles in CA1 cells were likely undergoing disengagement which could have also an impact on spindle pole integrity that leads to the formation of multipolar spindles independent of centrosome amplification – which is also an additional pathway for CIN.

Microtubule nucleation, the *de novo* formation of microtubules, is fundamental for the organization of microtubule arrays in both proliferating and postmitotic, differentiated cells^44^. Centrosomes represent the major microtubule organizing centers for nucleating microtubules in mammalian cells^83^, and some studies demonstrated that tumor cells with elevated microtubule nucleation from amplified centrosomes presented an enhanced invasiveness^46,57,84^. In our study, we hypothesized that cells with centrosome amplification would have increased nucleation capacity, especially for CA1 cells – which is the most invasive step in the progression. To shed light on the centrosome nucleation in BC progression, a microtubule regrowth assay was performed (Figure 1F and G). As depicted in the graph (Figure 1G), although there was no significant difference, the mean nucleation capacity for MCF10A cells was lower when compared to AT1, DCIS and CA1 cells (in foci with < 2 centrioles). For foci of 2 centrioles, there was no major difference among steps of BC progression. However, for foci with > 2 centrioles, we found a decrease in the nucleation capacity according to the progression of the disease, in contrast to what has been previously reported^46,57^. Our results show that microtubule nucleation capacity decreases according to disease progression in cells with centrosome amplification.

Taken together these results show that MCF10A series presented centriole loss and amplification and these characteristics increased with disease progression. Moreover, cells also presented a variety of mechanisms to cope with multipolar spindles.

### There is a loss of primary cilia (PC) according to BC progression and the frequency of PC increases according to the time of serum starvation in MCF10A

Centrioles, the core of centrosomes, are also essential for the formation of PC. PC is an organelle that protrudes from the cell surface into the extracellular space and functions as spatially restricted hubs, contributing to the regulation of cell signaling pathways^85,86^. Mouse models of PC demonstrate that, depending on the environment, the presence of cilia or their loss are required for tumor growth, playing a role in the tumorigenesis^87,88^. Thus, we analyzed the incidence of PC in MCF10A series (Figure 2A – C) after 5, 7, and 10 days of serum starvation using IF. Figure 2B shows that the percentage of cells with PC in MCF10A cells increased progressively with the duration of serum starvation. Moreover, the same trend was observed in RPE1 cells. The incidence of PC in AT1 cells was very low, with a mean percentage of 0.61% only at the day 7 of serum starvation. For the later stages (DCIS and CA1 cells), there was loss of PC, in disagreement with some studies which stated that PC can be found in those steps although in a low frequency^48,89,90^. Moreover, the size of PC was also analyzed (Figure 2C). For MCF10A cells, the mean size increased from day 5 to 7 and decreased at day 10. A similar trend was found in RPE1 cells as wellFurthermore, for AT1, the mean size was 7.6 at day 7 of serum starvation. Taken together, these results showed that there is a loss of PC according to BC progression and the frequency of PC increases according to the time of serum starvation in MCF10A.

**Figure 2.**
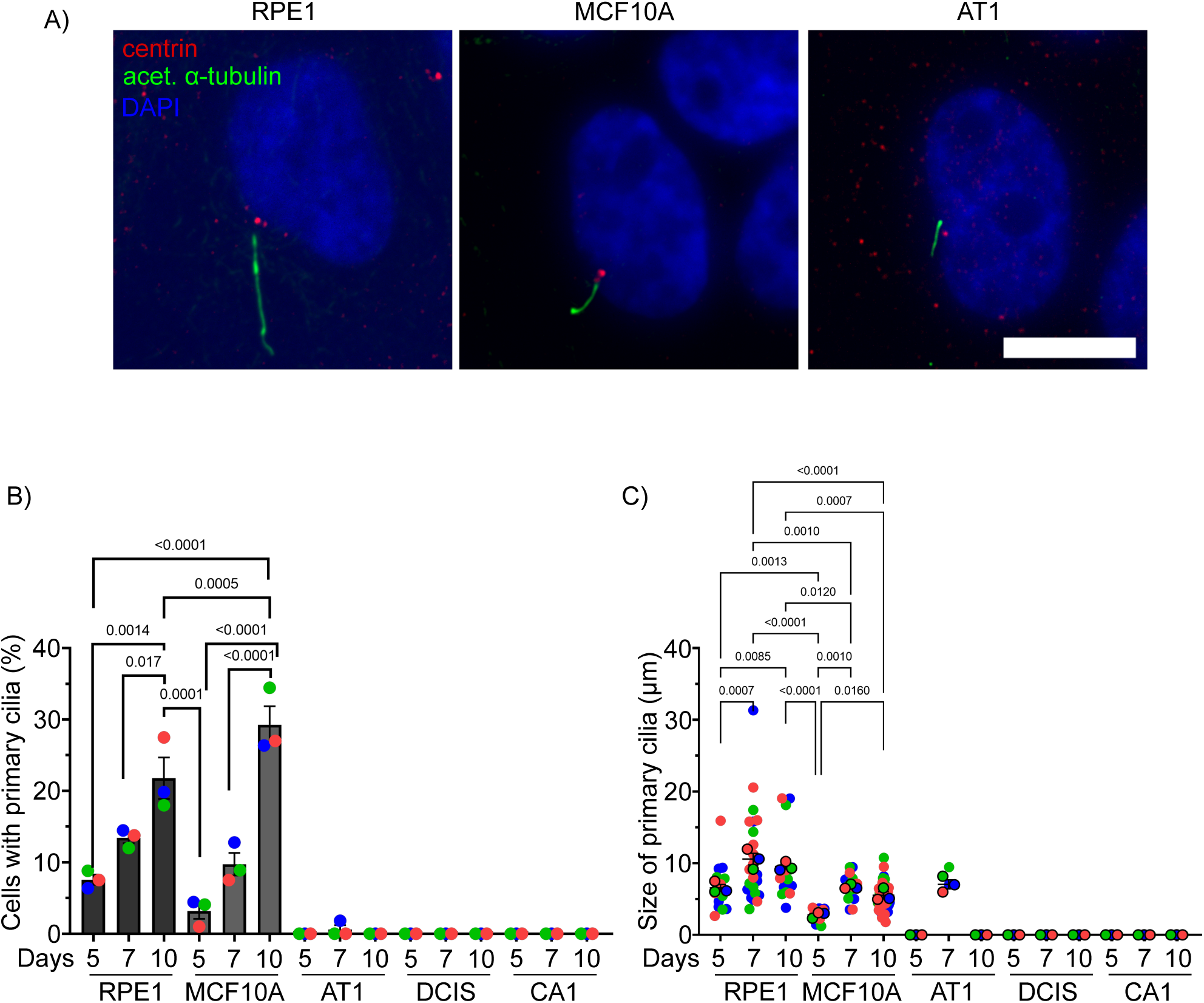
There is a loss of PC according to breast cancer progression and the frequency of PC increases according to the time of serum starvation in MCF10A. A) Representative image of cells with primary cilia stained with anti-centrin-2 (red), anti-acetylated α-tubulin (green) and DAPI. B) Graph shows the percentage of cells with primary cilia (PC) in MCF10A series and RPE1 (n≥= 350 cells). C) Size (in µm) of the primary cilia (PC) in MCF10 series and RPE1 (n≥= 20 cells). All statistical tests of comparative data were performed using one-way ANOVA for differential comparison between more than two groups. Data are shown as mean ± SEM. P < 0.05 were considered statistically significant. n = 3 biological replicates. Scale bar: 10µm

### Time spent in mitosis increases with disease progression probably due to errors of chromosome alignment

In order to examine the chromosome segregation in MCF10A series, live cell imaging was used to compare chromosomally stable cell lines (MCF10A^91^ and RPE1) and aneuploid/CIN cell lines of the series (AT1, DCIS, and CA1).

Firstly, we evaluated mitotic progression kinetics in the series in order to determine whether the control of mitosis was perturbed (Figure 3A – E). In the Figure 3A, the timeline of the experiment is shown (see material and methods for details). The frames showing the different mitotic stages measured are depicted in Figure 3B. As the disease progresses, we observed a significant prolongation of the time from nuclear envelope breakdown (NEB) to anaphase onset (AO) (Figure 3C). CA1 cells had the highest time spent in mitosis (42.28 min, p = 0.0025) followed by DCIS cells (38.67 min, p = 0.0183) when compared to MCF10A cells (28.15 min). Moreover, AT1 cells (30.98 min) had similar time when compared to MCF10A and RPE1 cells (30.52 min). Next, we determined the duration of the different mitotic phases. There was no difference in time from NEB to metaphase in AT1 (15.83 min) and DCIS cells (15.79 min) when compared to MCF10A cells (15.12 min) (Figure 3D). However, cells from CA1 cells spent a longer time to align at the metaphase plate (19.98 min, p = 0.0389), when compared to MCF10A cells (15.12 min). This likely happens because CIN cancer cells have excessive chromosome numbers and, in order to achieve stable attachments for all chromosomes, these cells will take longer time to align at the metaphase plate when compared to bipolar diploid cells^92^. Furthermore, CA1 (22.29 min, p = 0.0377) and DCIS cells (22.89 min, p = 0.0267) spent a longer time from metaphase to AO when compared to MCF10A cells (13.03 min) (Figure 3E). Furthermore, although AT1 cells (15.16 min) exhibited a longer metaphase to AO period, there was no statistical difference when compared to MCF10A and RPE1 (13.36 min). Collectively, these results showed that the time spent in mitosis increases with disease progression probably due to errors of chromosome alignment.

**Figure 3.**
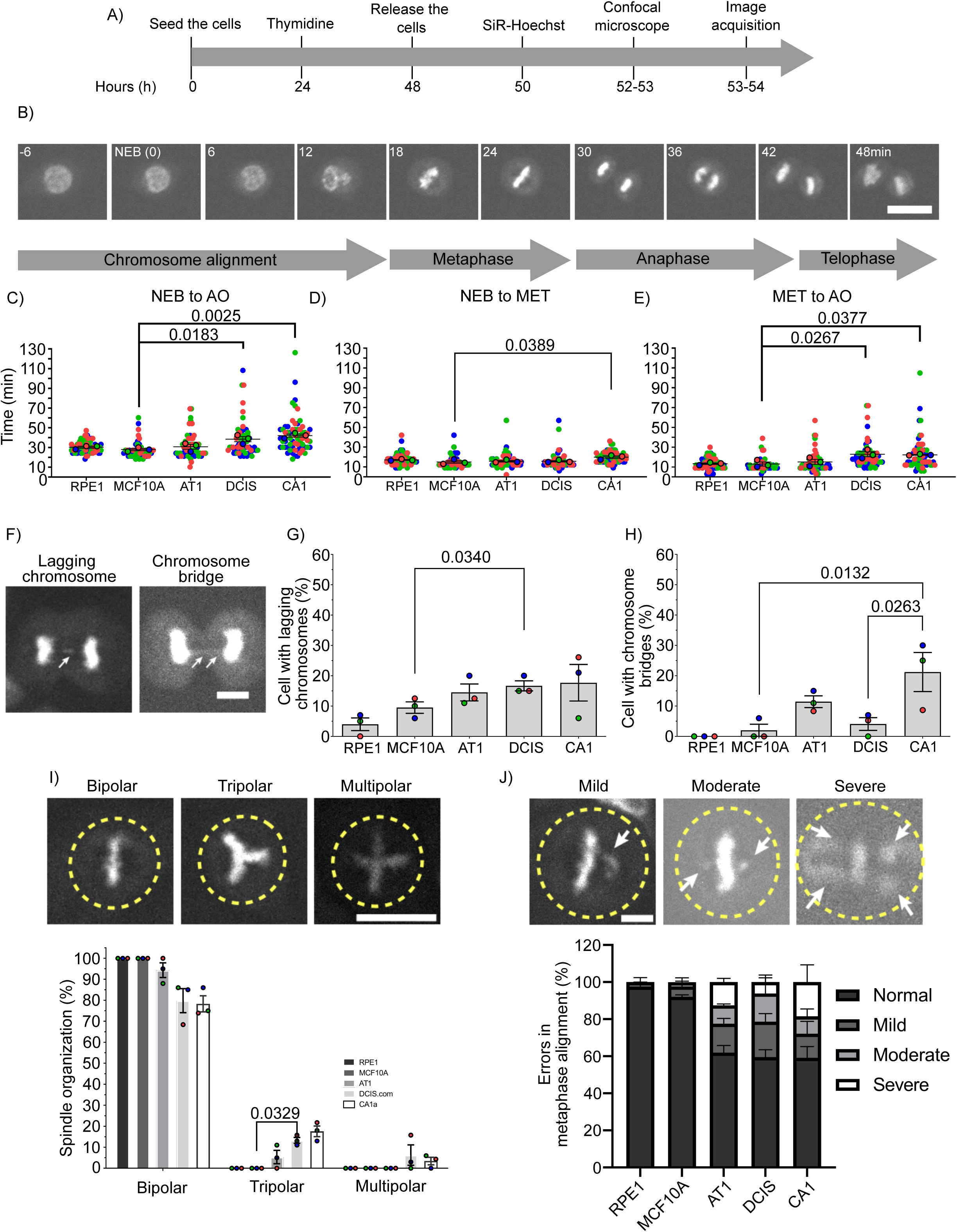
Time spent in mitosis increases with disease progression probably due to errors of chromosome alignment. A) Scheme of the experiment of live cell imaging. B) Representative time-lapse images of MCF10A stained with siR-DNA, showing the different phases of mitosis. C) Time spent in mitosis from NEB (nuclear envelope breakdown) to anaphase onset (AO). D) NEB to metaphase (MET). E) Metaphase (MET) to Anaphase onset (AO). F) Representative images of lagging chromosomes (arrow) (image on the left) and chromosome bridge (arrow) (image on the right) stained by siR-DNA. G) Graph shows the percentage of cells presenting lagging chromosome at anaphase. H) Graph shows the percentage of cells with chromosome bridges. I) Representative images of different spindle organizations – bipolar (left), tripolar (middle), and multipolar (right) spindles. Graph shows the percentage of cells presenting different spindle organizations. J) Errors in chromosome alignment in MCF10A series analyzed by live cell image. Representative image of different cells showing errors of chromosome alignment at metaphase: mild (1 chromosome), moderate (2 chromosomes) and severe (≥ 3 chromosomes). Graph showing the percentage of cells presenting mild, moderate and severe errors in chromosome alignment in MCF10A series. All statistical tests of comparative data were performed using one-way ANOVA for differential comparison between more than two groups. Data are shown as mean ± SEM. P < 0.05 were considered statistically significant. n = 3 biological replicates. Scale bar: 10µm (Figures A, D) and 5um (F and I).

Chromosome segregation defects are often detected in cancer cells, and are associated with poor tumor prognosis and overall survival in cancer^14^. Lagging chromosomes for example are the consequence of merotelic attachments (when a single kinetochore is attached to microtubules emanating from opposing spindle poles)^94,95^. Merotelic attachments are sufficient to silence the mitotic checkpoint^5^ and the resulting lagging chromosomes may lead to the formation of MNi^96–98^. Moreover, chromosome bridges consisting of chromatin connecting the two segregating masses of chromosomes and have been associated with CIN in human tumours^99^. These structures can be segregated into the a daughter nucleus or to form extranuclear MNi^100^. To study these defects in the MCF10A continuum series, we assessed the frequency of lagging chromosomes in anaphase and chromosome bridges in the series (Figures 3F – H). For lagging chromosomes in anaphase (Figure 3G), the percentage of cells increased according to disease progression. For the chromosome bridges (Figure 3H), the highest percentage was found in CA1 cells when compared to MCF10A cells. The percentage of cells presenting chromosome bridges in AT1 (11.04%) and DCIS cells (4.08%) was higher when compared to MCF10A cells (2%), although not statistically significant. There was no difference between MCF10A and RPE1 cells, as anticipated. Overall, these results show that the levels of these CIN markers (such as lagging chromosomes and chromosome bridges) increase across the disease. Moreover, these markers were present early in the continuum (AT1 cells) and reach high levels in the invasive cell line (CA1 cells), when compared to the chromosomally stable cell line (MCF10A cells).

Spindle multipolarity is a potential cause of asymmetric chromosome segregation and aneuploidy^55,81,92^.To determine whether cell lines from the MCF10A continuum present multipolar spindles, we analyzed spindle pole number in the series, scoring the cells with bipolar (2 poles), tripolar (3 poles) and multipolar (> 3 poles) spindles (Figure 3I). The percentage of cells presenting bipolar spindle decreased with the disease progression although there was no statistical significance among cell lines of the continuum and only bipolar spindles were identified in MCF10A and RPE1 cells (100% each). Furthermore, the percentage of tripolar spindle cells increased according to disease progression. Finally, cells with multipolar spindles (> 4 poles) were found in a very low percentage, only in DCIS and CA1 cells (6.26 and 3.44% of cells, respectively). These results showed that the aneuploidy/CIN cell lines (AT1, DCIS, and CA1 cells) are more prone to form multipolar spindles when compared to chromosomally stable cell line (MCF10A cells) in BC.

Sister chromatids that attach to the mitotic spindle but do not bi-orient (i.e. sister kinetochores that are not attached to microtubules extending from opposite spindle poles), or chromatids that lack spindle attachments altogether, are at risk for missegregation^101^. Erroneous chromosome congression and alignment leads to mitotic arrest or delay enforced by the spindle assembly checkpoint^102,103^. If that misalignment is not corrected, it will lead to the formation of aneuploid progeny and tumorigenesis^104–106^. To explore these errors in the context of the MCF10A continuum, we evaluated the frequency of errors in chromosome segregation during metaphase in MCF10A (Figures 3J). Most of the cells in MCF10A (93.28%) underwent a normal mitosis where all chromosomes aligned at the metaphase plate correctly. AT1, DCIS and CA1 cells presented high frequencies of cells with defects of chromosome alignment (37.94%; 35.68%; and 40.71%, respectively) when compared to MCF10A cells, with increasing severity as disease progresses. Overalls, these results show that most of the cells from the chromosomally stable cell lines (MCF10A and RPE1 cells) underwent normal mitosis, as expected, with few segregation errors, and CIN cell lines (AT1, DCI, and CA1 cells) presented cells with high frequency of misaligned chromosomes, potentially explaining the prolonged mitotic duration in these lines.

### Chromosome segregation defects increase with disease progression

Morphological indicators of CIN, including MNi, NBs, and NPBs help in prognostication of breast carcinoma^96,107^. To explore the presence of these structures in BC, we analyzed the frequency of cells presenting those morphological indicators by immunofluorescence (Figure 4A – E). The frequency of MNi in MCF10A series is depicted in the Figure 4B. The percentage of cells presenting 1 MNi was greater in AT1, DCIS and CA1 cells than in MCF10A cells, with progressively increasing frequency. The same trend was found in cells presenting 2 and 3MNi – with CA1 cells exhibiting the highest percentage of MNis in the series. Moreover, cells with 4 MNi were found only in CA1 cells. Similarly, the cells in the MCF10A continuum presented increasing percentage of NBs (Figure 4C), nucleoplasmic bridges (Figure 4D) and anaphase lagging chromosomes (Figure 4E). Interestingly, the majority of lagging chromosomes found were KT positive, a consequence of merothelic attachment (Figure 4E).

**Figure 4.**
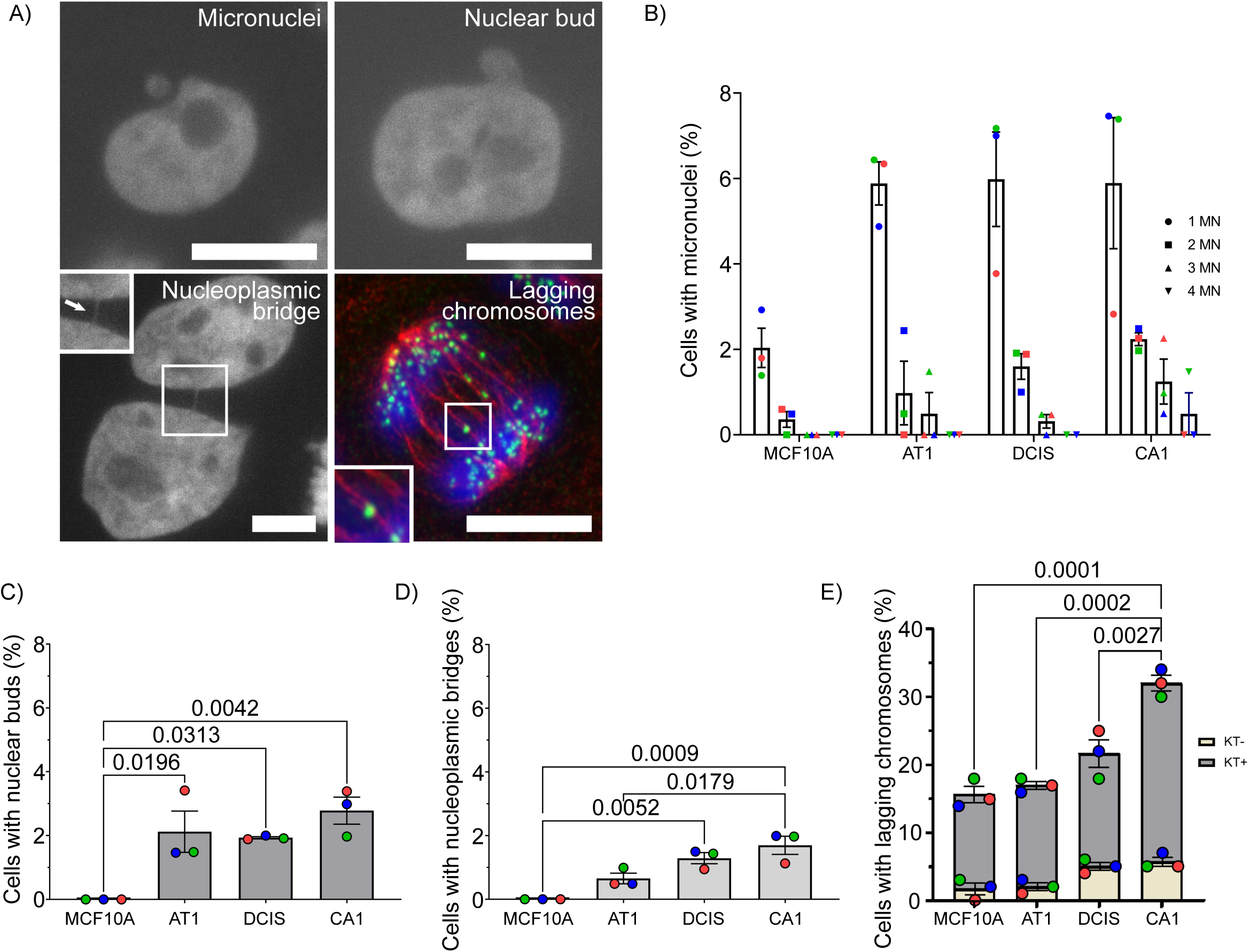
Chromosome segregation defects increase with disease progression. A) Mitotic defects in MCF10A series analyzed by immunofluorescence. Representative images of a micronuclei, nuclear bud, and nucleoplasmic bridges (inset shows a closer look of the bridge), labeled with DAPI. Immunofluorescence showing a lagging chromosome at anaphase labeled with anti-CREST (green), anti-α-tubulin (red), and DAPI (blue) (image at lower right). Inset shows a merotelic attachment (one kinetochore is attached to mitotic spindle from both spindle poles). Graphs showing the percentage of cells presenting micronuclei (B), nuclear buds (C), nucleoplasmic bridges (D), and lagging chromosomes at anaphase (E) in MCF10A series. All statistical tests of comparative data were performed using one-way ANOVA for differential comparison between more than two groups. Data are shown as mean ± SEM. P < 0.05 were considered statistically significant. n = 3 biological replicates. Scale bar: 10µm

Taken together, these results show that chromosome missegregation errors (including MNi, NBs, NPBs, and chromosome lagging, which are closely linked to CIN) increase with the disease progression in MCF10A series. Moreover, our results demonstrated that most lagging chromosomes were caused by erroneous kt-MT attachment, which could explain the increased frequency of MNi across the series.

### The overall kt-MT interaction is perturbed in the MCF10A series

kt-MT dynamics and attachment stability are finely regulated in order to prevent CIN, and kt-MT attachment dynamics is often deregulated in tumor cells^2,108^. Successful attachment between MTs and kinetochores enables sufficient force to be delivered through K-fibers (bundles of 20 – 40 kt-MTs attached to the chromosomes) to allow segregation of a chromosome into two daughter cells^109^. Thus, understanding k-fiber formation and dynamics is crucial for the study of mechanisms essential for chromosome alignment and segregation. To gain insights into how kinetochore–microtubule attachment stability aligns with the progression of BC, we subjected cells from the MCF10A series to cold treatment (Figure 5A and B). In this assay, when cells are cold treated, the less stable microtubules are depolymerized initially, leaving preferentially stabilized microtubule populations such as k-fibers (bundles formed by kt-MT) intact^110,111^. In Figure 5A, the timeline of the experiment is shown. Representative images of the experiment are also shown in the Figure 5B. The quantification of the experiment is depicted in Figure 5B (upper and lower graphs correspond to total tubulin intensity and tubulin intensity corrected by time 0, respectively). For the normal breast MCF10A cells, cold treatment produced the expected progressive destabilization of MTs, (see graph for tubulin corrected by time 0). After 20 and 30 min on ice, less than 35% were cold stable kt-MT. In contrast, in AT1, DCIS, and CA1 cells, cold treatment did not destabilize as much as in MCF10A cells. In fact, AT1, DCIS and CA1 cells presented higher levels of cold stable kt-MT bundles compared with the MCF10A cells. In some time points (30 min for AT1 and 20 min for CA1, for example), kt-MT attachments were even stronger (hyperstable) when compared to their respective control at time 0. Furthermore, centriole amplification did not have an impact on kt-MT stability (Supplementary Figure 1). The higher kt-MT stability found in AT1, DCIS, and CA1 cells may also explain the high frequency of missegregation defects in these CIN cell lines (Figures 3 and 4). K-fiber stabilization is also achieved through the Ran-importin beta-regulated protein HURP^112^. So we decided to quantify the expression of HURP in the MCF10A series (Figure 5C). MCF10A cells presented the highest HURP intensity when compared to AT1, DCIS, and CA1 cells. This accumulation of HURP in MCF10A cells can be correlated with higher intensity of k-fibers in MCF10A under basal, non-cold-treated conditions, when compared to AT1, DCIS, and CA1 cells (Figure Supplementary Figure 2). Taken together, these results showed that overall regulation of kt-MT interactions is disturbed in the MCF10A series.

**Figure 5.**
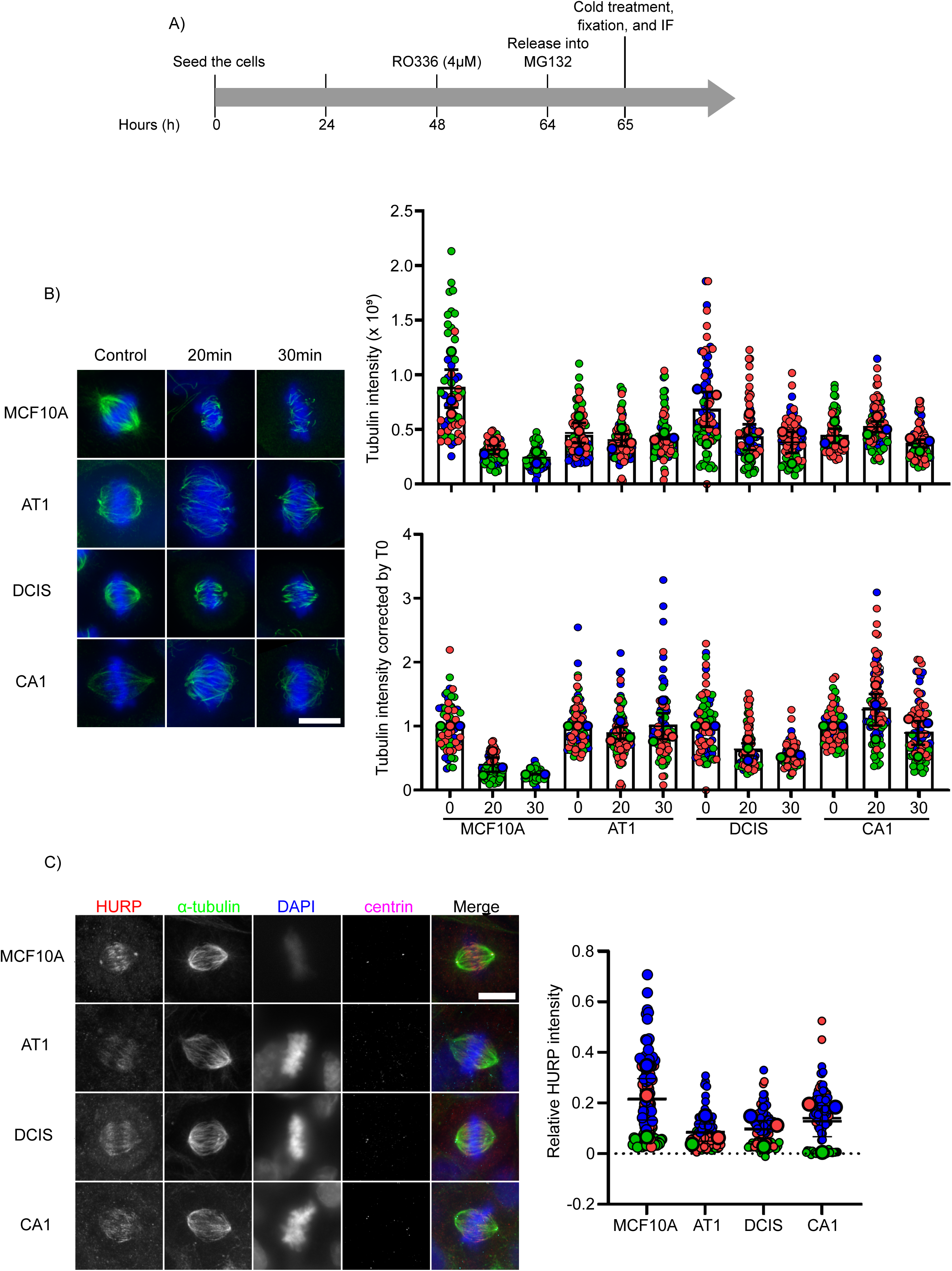
kt-MT interaction is more stable in benign, pre-malignant and invasive cells when compared to normal cell. A) Timeline of the cold stable assay. B) Representative images of MCF10A series non treated (control) and after 20 and 30 minutes of ice cold treatment. Cells were stained with anti-α-tubulin (green) and DAPI (blue). Quantification of MT density: upper graph corresponds to the total tubulin intensity and lower graph corresponds to tubulin intensity normalized by time 0 (T0). C) HURP expression in MCF10A series. Representative images of MCF10A series labeled with anti-HURP (red), anti-α-tubulin (green) and DAPI (blue). Graph shows the quantification of relative HURP intensity in MCF10A series. All statistical tests of comparative data were performed using one-way ANOVA for differential comparison between more than two groups. Data are shown as mean ± SEM. P < 0.05 were considered statistically significant. n = 3 biological replicates. Scale bar: 10µm.

### Centriole number has a major impact on the mitotic spindle orientation in MF10A series

Centrosomes serve as major microtubule-organizing centers (MTOCs) in animal cells and therefore play key roles in organizing spindle formation, chromosome segregation and cytokinesis but also polarity and motility^113^. We hypothesize that, if centrioles play an important role in organizing the mitotic spindle, any alteration in the centriole number would have an impact on spindle morphometrics. To analyze this assumption, morphometric analysis (including spindle length, half spindle length, chromosome congression, and spindle angle) (Figure 6A) was assessed in the BC progression using IF. Centriole number did not have an overall impact on spindle length for each cell line (Figure 6B). However, comparing cell lines with 2:2 centrioles, there was a statistical difference in spindle length where AT1 and CA1 cell lines (with 2:2 centrioles) presented a higher spindle length mean (11.77 ± 1.23, p = 0.0299, and 11.87 ± 1.24 µm, p = 0.0237, respectively), when compared to MCF10A cells(8.30 ± 1.26 µm). No significative alteration was found when comparing DCIS (10.30 ± 1.12 µm) with MCF10A. Furthermore, when comparing centriole loss or centriole amplification among different cell lines, spindle length mean did not change across the disease.

**Figure 6.**
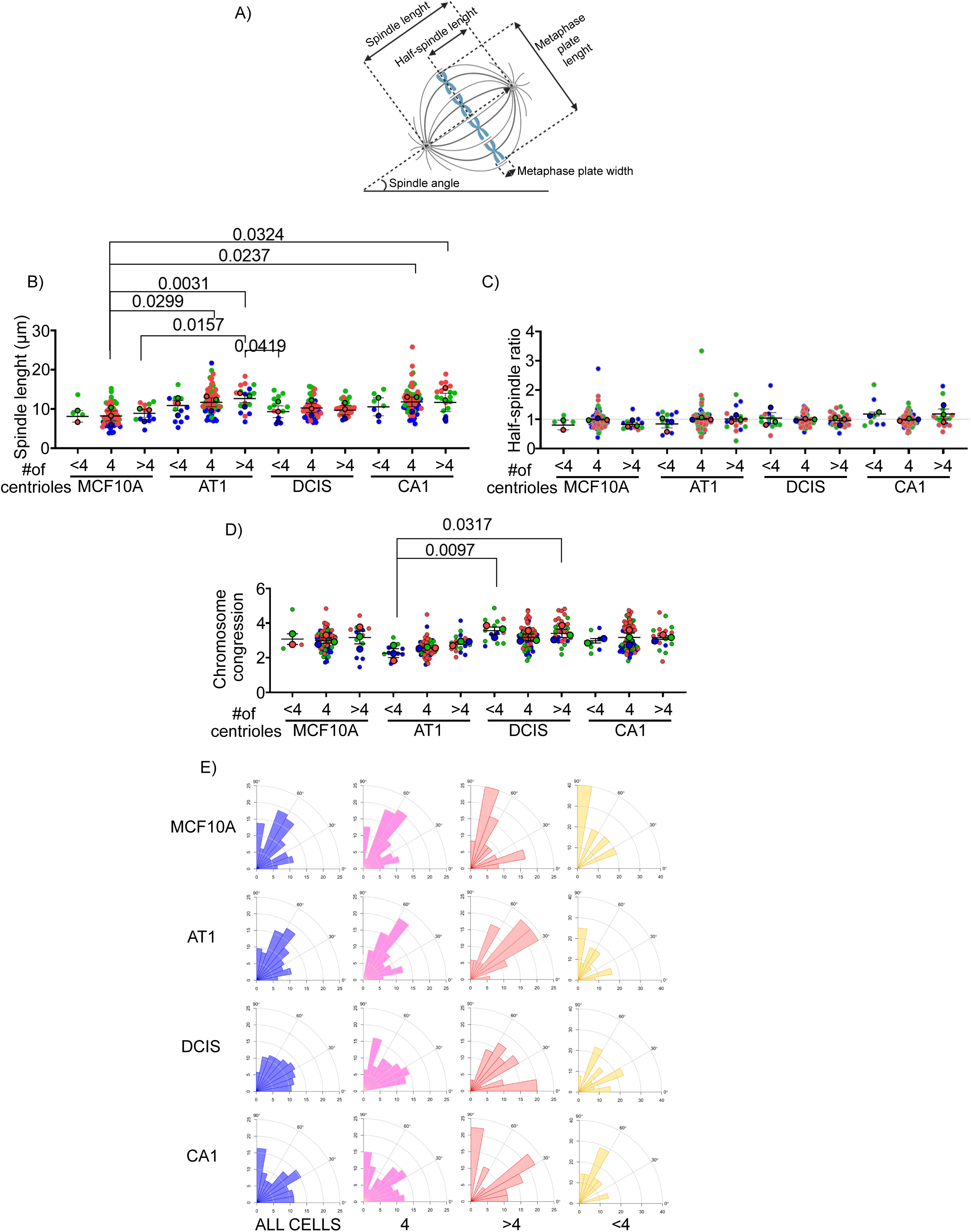
Centriole number has a major impact on the mitotic spindle orientation in MF10A series. Alterations in the mitotic spindle from MCF10A series. A) Representation of a mitotic spindle and the measurements analyzed. Spindle length corresponds to the distance from one pole to another. Half-spindle ratio is the ratio of distance from the poles till the middle of mitotic spindle. Chromosome congression is calculated based on the ratio of length and width of metaphase plate. Spindle angle was calculated between the pole– pole axis and the substrate plane. Superplots for the quantification of a spindle length (B), half-spindle ratio (C), and chromosome congression (D) in MCF10A series. E) Angular histograms for the mitotic spindle in MCF10A series. All statistical tests of comparative data were performed using one-way ANOVA for differential comparison between more than two groups. Data are shown as mean ± SEM. P < 0.05 were considered statistically significant. n = 3 biological replicates. <4: cells with bipolar spindle with less than 4 centrioles (centriole loss); 4: >4: cells with bipolar spindle with 2 centrioles per pole (normal cells); >4: cells with bipolar spindle with more than 4 centrioles (centriole amplification).

Next, half spindle ratio, which shows the position of the metaphase plate^50^, was also evaluated in the MCF10A series (Figure 6C). Values closer to 1 correspond to an equatorial plate position. When values deviate from 1, the metaphase plate is off centered which will lead to the formation of asymmetrical cell division which can lead to aneuploidy^50,114^. Comparing the number of centrioles (centriole loss, 2:2, or centriole amplification) inter and intra cell lines, the ratio was closer to 1 which represents nearly equal half-spindle lengths (Figure 5C). This result shows that centriole number does not have an impact on the position of the metaphase plate in this continuum, which means that even presenting centriole amplification, cells will present an equatorial plate position and will have symmetrical cell division.

Furthermore, in order to determine if the centrosome number affected the chromosome congression, we measured the ratio of the width to the height of the chromosomal mass (congression index) in the cells that reached the metaphase plate (Figure 6D). The number of centrioles (centriole loss, 2:2, or centriole amplification) did not change the chromosome congression, even comparing inter and intra cell lines – which shows that the chromosome congression is independent of the centriole number in this series.

Proper orientation of the mitotic spindle is critical and it determines the cell fate (symmetric versus asymmetric cells) and also dictates tissue architecture. Epithelial cells (like from human breast, for example) divide symmetrically to produce identical daughters^115,116^. Thus, failures in spindle orientation could cause loss of tissue organization which can promote tumorigenesis^117,118^. To explore this in the context of the MCF10A continuum, spindle angle was assessed as a readout for spindle orientation. In Figure 6E, the distribution of spindle angles in MCF10A series according to the centriole number is shown. For normal cells (4 centrioles), the distribution of spindle angle was similar between MCF10A and AT1 cells, and between DCIS and CA1 cells. Furthermore, when comparing cells with centriole amplification (>4 centrioles), the distribution of spindle angle changed completely across the disease – the same pattern was seen for cells with centriole loss (cells with <4 centrioles) despite the small number of cells. These results demonstrated that alteration in the centrosome number disrupted proper spindle orientation in MF10A series, which could lead to disruption of tissue architecture, a defining characteristic of cancer.

### HTA identifies negative regulators of ciliogenesis and centrosome biogenesis in MCF10A series

In order to explore the cause of the ciliogenesis and centrosome defects and identify regulators that could explain, for example, the loss of PC and some errors of chromosome segregation closely linked to CIN, such as lagging chromosomes in anaphase and centrosome amplification in the MCF10A series, we analyzed the HTA performed in MCF10A series already published by our group^38^. For this analysis, we set the absolute value of FC ≥ |2.0| and P < 0.05 as filter condition (Figure 7A). Figure 7B shows the Venn diagram depicting the numbers of overlapping and unique up and down regulated genes in the MCF10A series. In total, 2219 DEGs were identified, including 1165 genes (532 upregulated and 633 downregulated) in AT1, 1500 genes (759 upregulated and 741 downregulated) in DCIS, and 1473 genes (697 upregulated and 776 downregulated) in CA1. A comparison of these datasets found 338 and 337 overlapping DEGs across upregulated and downregulated genes (Figure 7B).

**Figure 7.**
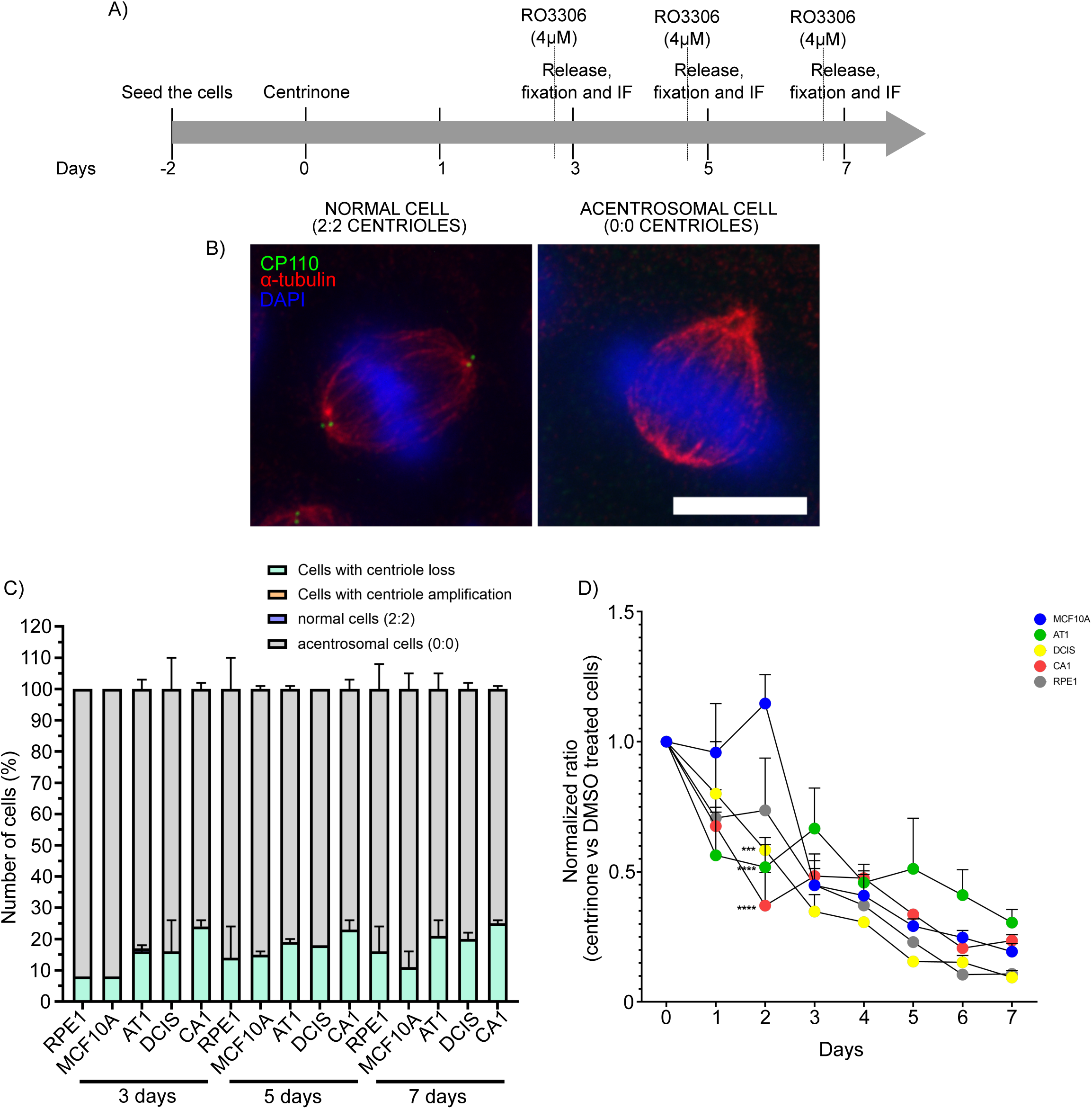
AT1 and CA1 proliferate more in the absence of centrosomes. A) Timeline of the experiment. B) Representative images of a normal and acentrosomal cell, non treated (DMSO) and treated with centrinone (200nM), respectively. Cells were stained with anti-α-tubulin (red), anti-CP110 (green), and DAPI (blue). C) Graphs showing centrosome number distribution in MCF10A series after 3, 5 and 7 days under centrinone treatment (200nM). D) Proliferation curves of MCF10A series with or without centrinone treatment (200nM) during 7 days. Statistical tests of comparative proliferation data were performed using one-way or ANOVA for differential comparison between more than two groups. Data are shown as mean ± SEM. P < 0.05 were considered statistically significant. n = 3 biological replicates. Scale bar: 10µm.

**Figure 8.**
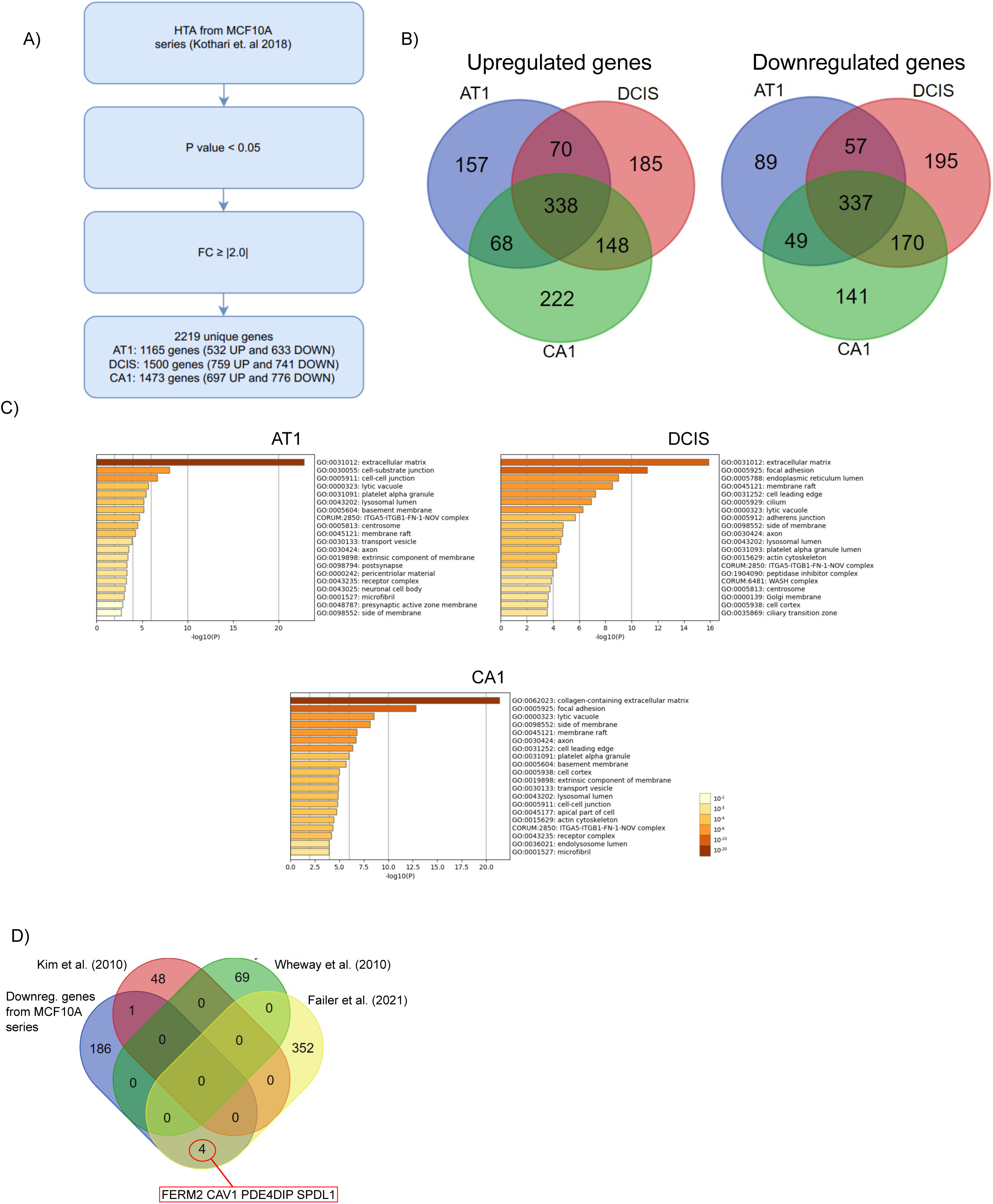
HTA identifies negative regulators of ciliogenesis and centrosome biogenesis in MCF10A series. A) Flowchart of study design. HTA: human transcriptome array. FC: fold change. B) Venn diagram showing the DEGS (up- and downregulated genes) in each cell line (AT1, DCIS, CA1) compared to normal BC cell line (MCF10A). C) Enrichment analysis (Cellular components) of differentially expressed genes (DEGs) in MCF10A series using Metascape. Bar graph of enriched terms across input gene list, colored by *p*-values. G) Venn diagram comparing the 191 downregulated gene list that may play a role in ciliogenesis and centrosome duplication/biogenesis with 3 different gene lists from studies that used high-throughput siRNA screens to identify regulators of ciliogenesis.

For the functional annotation of DEGs, Metascape was used (Figure 7C and Supplementary Figure 3). The main enriched GO terms were categorized based on pathways and cellular components to identify characteristics that could be related to cilia, mitotic spindle, and centrosome biogenesis. The major pathways in upregulated genes from AT1, DCIS and CA1 were associated with inflammatory processes (for example, hallmark TNFA signaling via NFKB, hallmark KRAS signaling up, hallmark inflammatory response), organogenesis (for example tube morphogenesis, tissue morphogenesis, cell morphogenesis), and hormone stimulus (hallmark estrogen response early, for example) (Supplementary Figure 3). Downregulated genes from AT1, DCIS, and CA1 were mostly enriched in pathways related to epithelial mesenchymal transition (ECM) (hallmark epithelial mesenchymal transition, for example), organogenesis (vasculature development, heart development, and cell morphogenesis, for example), and extracellular matrix organization (NABA core matrisome and extracellular matrix organization) (Supplementary Figure 3). Moreover, upregulated genes were enriched in cellular components related to cell-cell junctions, extracellular matrix, and membranes (for example, Golgi membrane, membrane raft, lyctic vacuole) (Supplementary Figure 3). Interestingly, downregulated genes were also enriched in GO cellular components related to cilia and centrosome (for example, pericentriolar material, centrosome, actin cytoskeleton, cilium) (Figure 7C). In order to gain a more detailed view about the GO terms involved in cilia, spindle and centrosome biogenesis, we compared the DEGs from AT1, DCIS, and CA1 and we created a heatmap with the top 100 GO terms from Cellular components (Supplementary Figure 4) using Metascape. The heatmap from downregulated genes showed more GO terms related to cilia and centrosome biogenesis (13 terms out of 100) when compared to upregulated genes (5 out 100). These results taken together showed that most of the phenotypes related to cilia, spindle and centrosome found in MCF10A series may likely be a consequence of downregulated gene expression.

Next, to gain insight about which downregulated gene(s)/pathway may play a role in the phenotypes found in MCF10A series, we compiled all the GO terms related to cilia, spindle and centrosome from the heatmap of Supplementary Figure 4 and we extracted all the genes. Table S1 shows the list of 191 downregulated genes found. From that list of genes, a PPI network and module analysis was performed in Metacape and visualized in Cytoscape to identify essential proteins at the network level. The Supplementary Figure 5 illustrates the PPI network from these 191 downregulated genes. The network found was highly noded and presented 149 nodes and 324 edges. Further analysis using the MCODE identified 8 functional modules in the PPI network. The top 4 functional modules were characterized as follows. The top one module (MCODE1) consisted of 7 genes, which included *PLEC*, *TGFB1I1*, *CLASP1*, *ATM*, *HOOK3*, *DGL5*, and *ABLIM3*. Most of these DEGS were enriched for GO terms related to focal adhesion, cell-substrate junction, and centrosome. Moreover, the top 2 module (MCODE2) consisted of 6 genes, which included *COL4A5, COL4A6, COL6A3, COL8A1, COL12A1*, and *P3H1*. These genes were mostly associated with GO terms related to collagen trimer, endoplasmic reticulum lumen, and collagen-containing extracellular matrix. Five genes comprised the MCODE3, including *MYL6, MYLK, TPM2, MYL9*, and *LRKK2*. The enrichment analysis for these were mostly related to actin cytoskeleton. Lastly, the top 4 (MCODE4) encompassed 4 genes (*KIFAP3, IFT81, IFT80, WDR19*) mostly enriched with GO terms related to ciliary tip, intraciliary transport particle, and cilium. Taken together, these results demonstrated the importance of these genes in processes related to centrosome, cilia, and actin cytoskeleton.

Cilia related proteins can also be found in non-cilia sites (i.e. sites outside the cilia) and may have non canonical functions, such as several mitotic events (for example, mitotic spindle assembly, mitotic spindle orientation, SAC, cytokinesis, etc.) (see review in Vertii and colleagues 2015^91^). AbouAlaiwi and colleagues (2011) demonstrated that cells with aberrant PC from endothelial cells presented multipolar spindles, centrosome amplification and no SAC activity, due to the downregulation of the chromosomal passenger complex survivin^119^. We thus wanted to test the hypothesis that downregulation of cilia proteins may play a role in the missegregation defects found in the MCF10A series. In order to determine negative regulators of the ciliogenesis, we compared our list of 191 genes with three different high-throughput siRNA screening studies designed to identify negative regulators of ciliogenesis in RPE1 cells^120^, mouse inner medullary collecting duct (IMCD3) cells^121^, and the Hs578T basal-like breast cancer cells (a cell line derived from a triple negative breast cancer patient)^122^. The Figure 7D illustrates the Venn diagram comparing our 191-downregulated gene list with the 3 siRNA screening studies, and shows poor overlap among the 4 studies, potentially as a result of different experimental conditions. Despite this, our gene list shared 4 genes (*FERMT2, CAV1, PDE4DIP*, and *SPDL1*) with the study from Failler and colleagues (2021), and 1 gene (*PARVA*) with the study from Kim et al (2010) that may be good candidates to explain the phenotype found in the MCF10A series. Interestingly, the 4 overlapping genes identified in both our study and the study from Failler et al. code for proteins with known mitosis/chromosome segregation functions. Studies show that kindlin-2 (product of the *FERMT2* gene) –depleted neuroblastoma SH-SY5Y cells exhibited delayed mitosis, spindle abnormalities and mitotic spindle assembly^124^. Caveolin-1 (*CAV1* gene product), which is the principal structural protein of caveolae, plays role in signaling transduction and intracellular trafficking of cellular components^125,126^. At the onset of mitosis, before chromosome segregation, Caveolin-1 is enriched at cortical regions and guides spindle orientation^127^. Spindle apparatus coiled-coil protein 1 (SPDL1) encodes a protein containing a helical domain which plays an important role in mitotic spindle formation and chromosome segregation^128,129^. KD depletion results in a long mitotic time, severe chromosome misalignment, and striking spindle rotation^130^. The *PDE4DIP* protein product anchors and sequesters components of the cAMP-dependent pathway to Golgi and/or centrosomes^131^. This protein forms a complex with AKAP9, EB1/MAPRE1 and CDK5RAP2, to promote centrosomal microtubule nucleation and extension, a pivotal process for cell migration and mitotic spindle orientation^132^.

Taken together these results showed that the downregulation of regulators of ciliogenesis may play a role in the missegregation defects found in the MCF10A series.

## DISCUSSION

The present study illustrated the importance of MCF10A series as a model for studying BC development. We used this BC continuum of isogenic cell lines, which included from normal breast to premalignant and malignant lessions with metastatic capabilities, and identified a range of mitotic defects that may promote the development and progression of BC. A characteristic of those mitotic errors (including lagging chromosome, NPBs, MNi, NBs, and centrosome amplification) found in MCF10A series is that they increased with the disease progression. Furthermore, our data demonstrated that these mitotic defects were already present in ADH, which means they can generate CIN/aneuploidy during the early stages of the tumorigenic process and perhaps contribute to the progression of early lesions to more aggressive cancers.

Centriole amplification was also evident in the series and most of the excess centrosomes in AT1, DCIS, and CA1 were generated by centriole overduplication. This mechanism indicates that the centriole overduplication was likely to be caused by a deregulation in the centrosome cycle. Interestingly, the HTA analysis identified many downregulated genes related to centriole that could explain the centriole overduplication. Furthermore, centriole disengagement was also found in the CA1 cells– which may also be related to an increased time spent in mitosis for that cell line. This alteration could represent an additional way of forming multipolar spindle independent of centrosome amplification, which will have an impact on spindle pole integrity and lead to missegregation of chromosomes and aneuploidy^81^.

Our study demonstrated that centrosome amplification likely disrupted mitotic spindle orientation in MCF10A series. This is an additional mechanism to generate chromosome missegregation, asymmetrical cell division and, therefore, aneuploidy. The correct orientation of the mitotic spindle is known to depend on the pulling forces acting between the spindle poles and cortical microtubule attachment sites, emanating from centrosomes^113^. Our results are in agreement with other study that showed centrosome amplification induced defects in spindle orientation^122,133^. However, centrosome amplification did not change other spindle morphometrics, such as spindle length, half-spindle ratio and chromosome congression, suggesting that these spindle morphometrics in MCF10A series are tightly regulated upon differences in centrosome number.

The structural integrity of the microtubule attachment sites on chromosomes is critical to provide a faithful chromosome segregation. Nevertheless, errors in the orientation of kt-MT attachments usually occur, especially in early phases of mitosis - but most of them are corrected^94^. One of these errors, named merotely, happens when a single kinetochore binds microtubules oriented toward both spindle poles^134^. The persistence of merotely leads to chromosome mis-segregation because merotelic kinetochores experience poleward force toward both spindle poles. Merotelic divisions can cause chromatids to lag behind, hindering their segregation to spindle poles^135,136^ Furthermore, tumor cells with CIN have elevated rates of merotelic attachments and lagging chromosomes^135,136^. In the present study, the kt-MT interaction is more stable in pre-malignant, malignant, and invasive cells when compared to normal cell in BC. This slow release from the kt-MT in the cancerous cell lines will increase the probability of mis-attachment and, therefore, whether this error persists, it will lead to the formation of lagging chromosomes in anaphase. Considering our results, we hypothesize that this may happen in the MCF10A series. Unstable cell lines (AT1, DCIS, and CA1) presented a higher kt-MT stability which leads to the formation of lagging chromosomes and MNi.

Our results showed that the presence of lagging chromosomes increased according to disease progression, and MCF10A cells had the lowest percentage followed by AT1, DCIS and CA1 cells. This raised a question whether, at later stages of disease progression, there was any mechanism of error correction compromised. Recent findings demonstrated the existence of a midzone-based Aurora B phosphorylation gradient that locally stabilizes KT-MT attachments by phosphorylating PLK1 at KTs, to promote efficient mechanical transduction of spindle forces at the kinetochores to correct errors in anaphase and prevent MNi formation during cell division^97,135,136^. However, additional experiments (IF to analyze the Aurora B gradient in anaphase, for example) are needed to evaluate whether AT1, DCIS, and CA1 cells present a compromised Aurora B gradient when compared to MCF10A cells. Furthermore, besides the presence of lagging chromosomes and chromosome bridges, another source of MNi was the presence of chromosome misalignment in MCF10A series (Figure 3J). Higher percentage of MNi (above 6%) in AT1, DCIS, and CA1 cells, may be also explained by presence of misalignments in these cell lines that eventually do not form lagging chromosomes. In agreement with our results, Gomes and colleagues (2022) showed that misaligned chromosomes that satisfy the SAC often directly missegregate (without lagging behind in anaphase) and have the highest probability to form micronuclei, specifically in human cancer cell models^137^.

Surprisingly, our results demonstrated that microtubule nucleation capacity decreases according to disease progression in cells with centrosome amplification (figure 1G). This finding is in contrast to reports that centriole amplification leads to an enhancing of MT nucleation and chromosomal instability (CIN)^46,66,84^. For example, in the context of BC, Lingle et al. revealed that microtubule-nucleation capacity is enhanced in BC cells with centriole amplification^66^. However, Kushner and colleagues found that supernumerary centrosomes in endothelial cells presented a reduced MT nucleation^144^ This difference in the result supports the view that the microtubule-nucleation ability of aberrant centrosomes might be either reduced or enhanced, depending on the identity and modification state of the overexpressed pericentriolar material components that surround the centrosome^76^.

Oncogenic Ras mutations, which can lead to the production of permanently activated Ras oncoprotein, occur in ∼25% of human cancers and regulate a complex signaling network that promote transformation^145^. Ras pathways regulate diverse cellular processes, including, cell proliferation, cell survival regulation, cytoskeleton rearrangement, membrane trafficking, cell death, cell motility^146^, nevertheless, little attention is given to its role on mitotic processes that ensure chromosome segregation fidelity^147,148^. Recently, Herman and colleagues (2022) proposed that the hyperactivation of HRas^G12V^ stimulates kinetochore kinases, including Aurora B, that leads to the hyperphosphorylation of kinetochores and decreased microtubule binding capacity, which compromises the chromosome alignment and segregation processes, and likely contributes to aneuploidy and/or chromosome instability^148^. Ganguli and colleagues (2023), after inducing the expression HRas^G12V^ in MCF10A, found that the oncogene HRas^G12V^ altered substrate adhesion during mitosis, resulting in misregulation of spindle orientation and post-mitotic respreading^149^. In our study, we can also hypothesizethe effect of active Ras in the chromosome segregation in MCF10A series, considering that AT1, DCIS, and CA1 cells harbor the activating mutation of HRAS (G12V)^30^ (Figure 1A). So, the higher percentage of chromosome errors and spindle orientation defects found in AT1, DCIS, and CA1 cells may also partially be explained by the Ras activation in those cell lines when compared to MCF10A.

We also investigated the presence of PC in the MCF10A series. Our results showed that there was a loss of PC according to BC progression and the frequency of PC increased according to the time of serum starvation in MCF10A series. We used our already published HTA performed in MCF10A series^38^ to discover the root of the ciliogenesis and centrosome defects and identify regulators that could explain the chromosome missegregation found in the continuum. Most of the centrosome and cilia related genes were downregulated in our dataset. Thus we hypothesized that the downregulation of cilia proteins may play a role in the missegregation defects found in the MCF10A series, considering some non canonical cilia functions related to mitotic spindle assembly, mitotic spindle orientation, SAC, and cytokinesis. Next, we created a list with 191 genes related to centrosome and cilia defects and compared with some siRNA screening studies, and found 4 candidate genes (*FERMT2*, *CAV1*, *PDE4DIP*, and *SPDL1*). Previous reports have described cilia proteins with cilia independent functions, playing role in mitotic spindle orientation, mitosis and mitosis checkpoint^150–152^. Remarkably, all the genes had functions in mitosis/chromosome segregation, which can suggest that they may play a role in the phenotype found in the MCF10A series. Nevertheless, further experiments (RT-qPCR to validate the genes and experiments of overexpression of the candidate genes) are needed to explore which gene is responsible for the phenotype found in the MCF10A series.

A simplified mechanism that explains how PC proteins can play a role in the missegregation defects is ilustrated in the Figure 9. PC formation and centriole biogenesis are highly regulated, and tightly linked to the cell cycle. In normal cells, upon cell cycle entry (G1), ciliary resorption/disassembly begins and the balance of cilium assembly and disassembly is shifted toward disassembly. Following cell division, at mitotic entry, the two centrosomes separate and move to the poles where they nucleate MTs to form the bipolar spindle. Then, cells will undergo correct chromosome alignment and originate euploid cells. In abnormal cell, downregulation of some genes impair/block the formation of primary cilia in G1. Then, following the cell division, it will lead to the formation of aberrant centrosomes^119^. Aberrant centrosome will undergo mitosis and move to the opposite poles. These centrosomes will form hyperstable kt-MT attachments which lead the chromosome missegregation errors that form lagging chromosomes. The resultant daughter cells will be aneuploid^153^.

**Figure 9.**
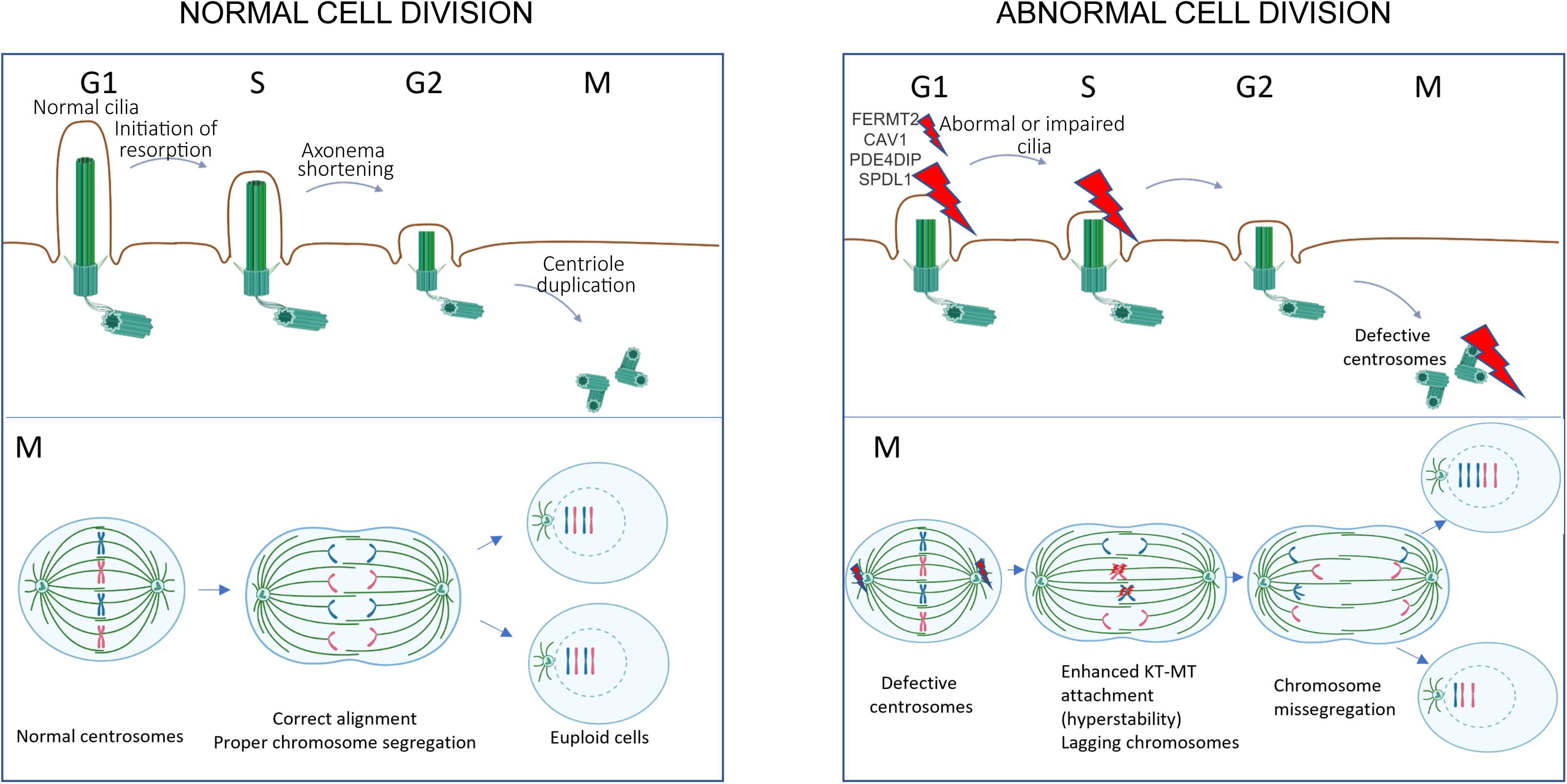
Schematic model indicating how the downregulation of PC can lead to the chromosome missegregation in MCF10A series.

## CONCLUSION

Using an *in vitro* model of disease progression (the MCF10A series of BC continuum), we characterized the missegregation defects that occur during the progression of the disease and lead to aneuploidy/CIN. MCF10A series presented several chromosome segregation defects, including lagging chromosomes, micronuclei, nuclear buds, nucleoplasmic bridges, errors of chromosome alignment, centrosome loss/amplification, which increased with the disease progression. Premalignant, malignant and invasive cell lines presented hyper stable KT-MT attachment, which can also explain the presence of errors of chromosome alignment in those cell lines. Furthermore, there is a loss of PC across the disease progression. From our HTA performed in the MCF10A series, we were able to find a group of genes related to cilia, centrosome and spindle and proposed that a downregulation of a cilia protein is responsible for the missegregation defects found in the series.

---

**Table 1.** List of 191 downregulated genes that may play a role in ciliogenesis and centrosome duplication/biogenesis and their respective fold change.

## Supplementary Figures

**Supplementary Figure 1.**
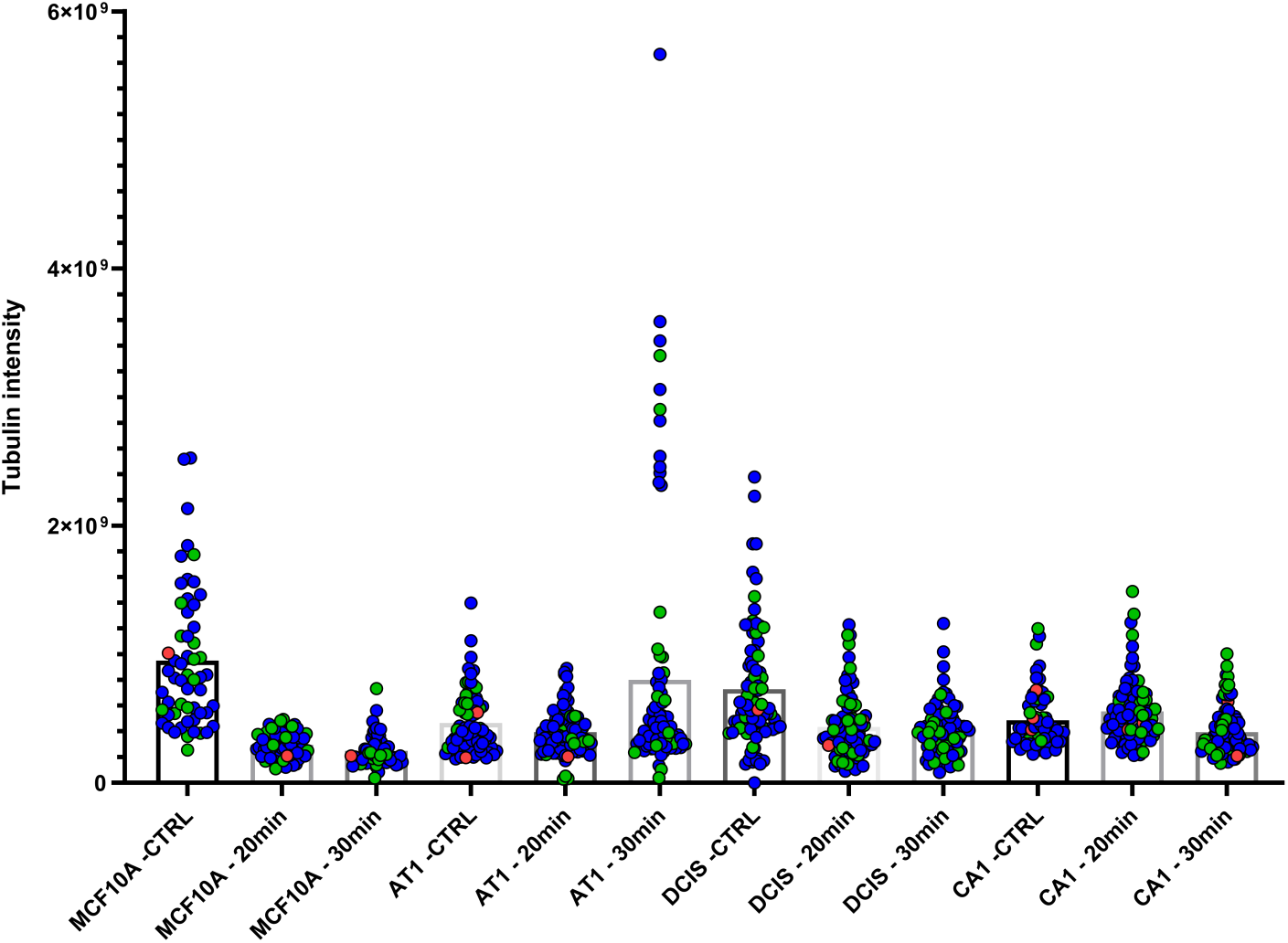
Quantification of MT density. Graph shows the total tubulin intensity in MCF10A series after 20 and 30 minutes of ice cold treatment. Blue dots: normal cells (4 centrioles); green dots: cells with centriole amplification (>4 centrioles); red dots: cells with centriole loss (<4 centrioles).

**Supplementary Figure 2.**
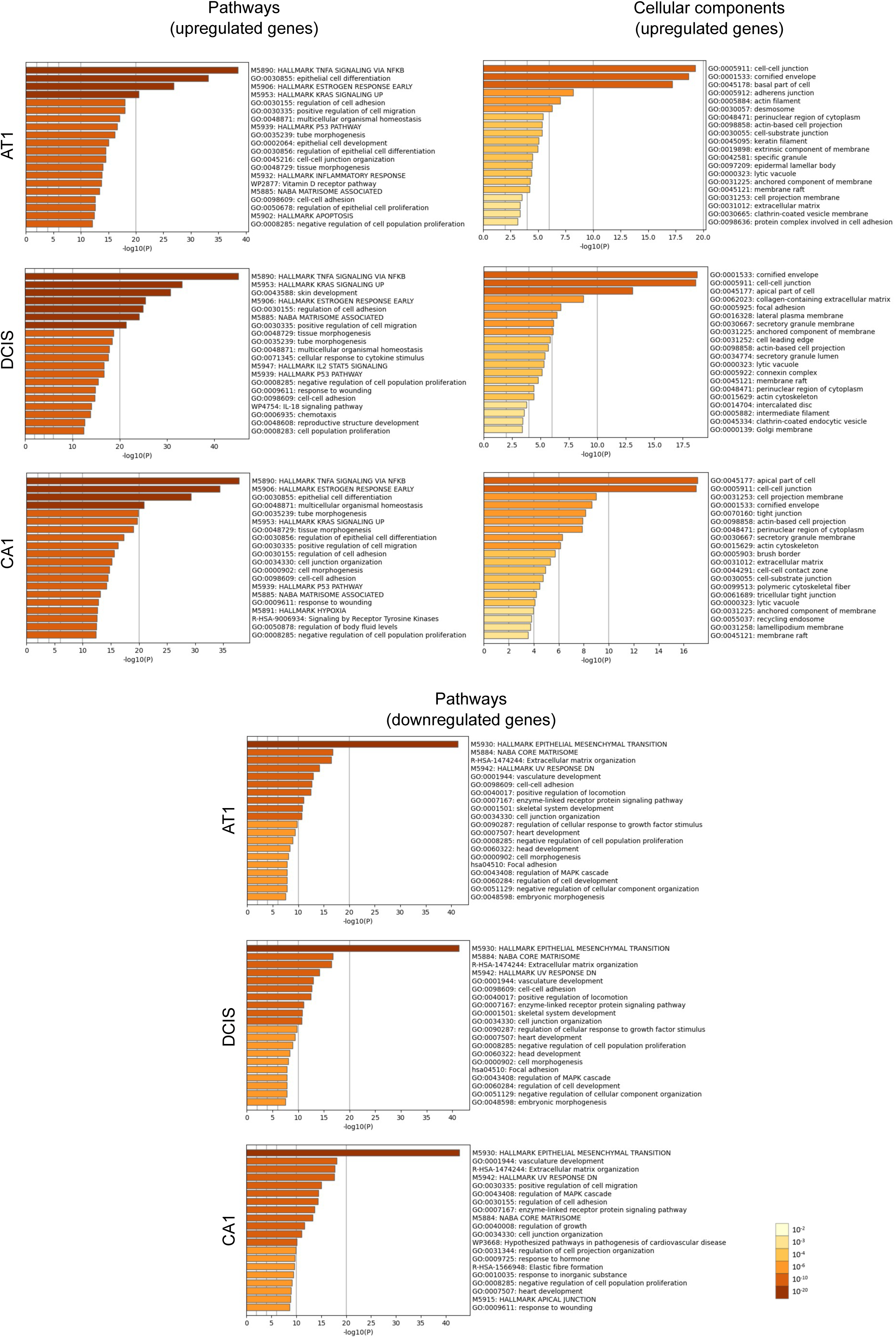
Enrichment analysis for upregulated genes (Pathways and Cellular components) and downregulated genes (Pathways) of differentially expressed genes (DEGs) in MCF10A series using Metascape. Bar graph of enriched terms across input gene list, colored by *p*-values.

**Supplementary Figure 3.**
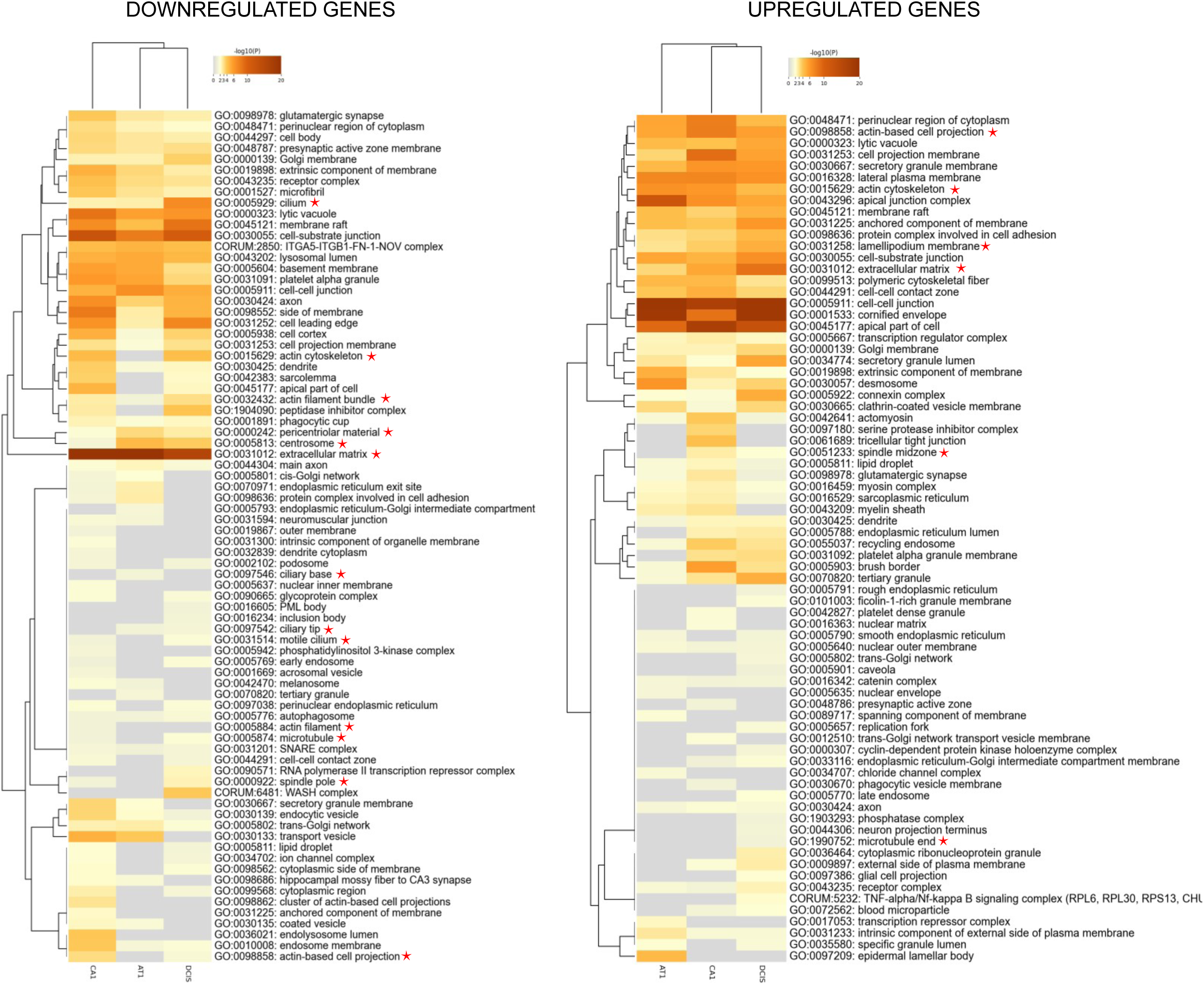
Enrichment Ontology cluster across down- (left side) and upregulated (right side) genes, colored by p-values, from MCF10A series (AT1, DCIS, and CA1). This heatmap corresponds to the top 100 clusters from Cellular components. Red stars correspond to gene ontologies related to cilia, centrosome and spindle.

**Supplementary Figure 4.**
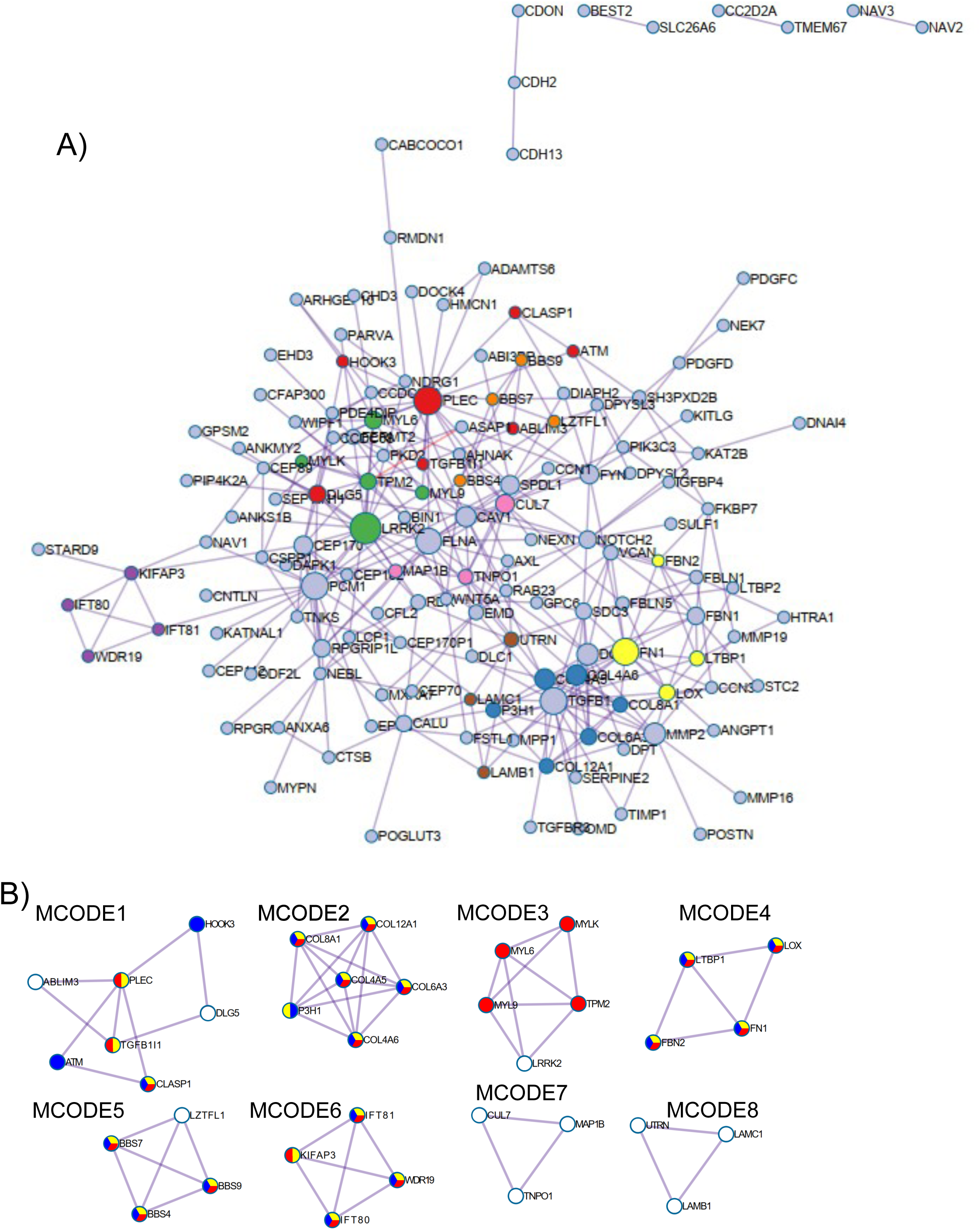
Protein-protein interaction network and MCODE components identified in the 191 downregulated gene lists.

